# Congruent neuronal modulation across competing actions challenges the role of the substantia nigra in action selection

**DOI:** 10.1101/2025.02.18.638935

**Authors:** Lingfeng Hou, Derin Timuçin, Olivia Sato, Sofia Grijalva Torres, Daniel Svedberg, John Assad

## Abstract

The basal ganglia are involved in the control of movement, but their exact role is unclear. Paradoxically, most of the inhibitory projection neurons in the main output nuclei increase firing around the time of movement; only a small fraction decrease firing. This antagonistic activity pattern could subserve action selection, with the small “decrease” population selectively disinhibiting the desired movement, and the larger “increase” population inhibiting competing movements. The action-selection hypothesis makes an implicit assumption: neurons that decrease firing to disinhibit a specific action should increase firing to inhibit that action when a different action is desired. To test this hypothesis, we recorded projection neurons in the substantia nigra pars reticulata (SNr) of mice trained to alternate between two different types of movements. Many SNr neurons showed a “ramping” pattern of pre-movement firing-rate modulation, with most neurons increasing firing, consistent with previous findings. However, contrary to the action-selection model, the overwhelming majority of SNr neurons exhibited congruent modulation between the competing actions, either increasing or decreasing their firing rates for both actions; only a small fraction of neurons exhibited opposite signs of modulation. Similar results were observed among SNr neurons identified as projecting to the arm or orofacial zones of motor superior colliculus. In addition, brief pauses in SNr firing outside explicit operant tasks were not accompanied by stereotyped movements. Our results are not easily reconciled with simple antagonistic mechanisms for action selection in the basal ganglia output nuclei. We also found that ramping activity in SNr neurons typically began hundreds of ms before self-timed and spontaneous movements, in contrast to previous findings suggesting that basal ganglia output is modulated too late to be involved in movement initiation. Our findings suggest constraints – and raise new questions – about the role of the basal ganglia in movement initiation and action selection.

## Introduction

The basal ganglia (BG) are a set of evolutionarily conserved sub-cortical nuclei involved in movement, motivation and other cognitive functions. The major input nucleus of BG, the striatum, receives extensive cortical and thalamic input, and in turn projects (directly and indirectly) to the major output nuclei of BG, the globus pallidus interna (GPi) and substantia nigra pars reticulata (SNr). The GPi and SNr contain tonically active GABA-ergic projection neurons which innervate thalamic and brainstem targets (Deniau et al., 2007; Nambu, 2007). Ascending projections to thalamic motor nuclei and descending projections to brainstem premotor nuclei in principle allow the basal ganglia to exert control over movement (Arber & Costa, 2022; Frost-Nylén et al., 2024; McElvain et al., 2021), but the mechanisms by which basal ganglia output influences movement remain unclear.

In the oculomotor system, the tonic discharge of SNr projection neurons provides continuous inhibition to the superior colliculus (SC) between saccades (Hikosaka & Wurtz, 1983a). At the initiation of saccades, many SNr neurons pause their tonic firing (Hikosaka & Wurtz, 1983b; Joseph & Boussaoud, 1985); this release of inhibition could allow a burst of collicular activity that triggers eye movement. In this view, basal ganglia output could gate movement via *disinhibition* (Deniau & Chevalier, 1992; Hikosaka & Wurtz, 1983c). More recently, transient dips in activity have been reported in mouse SNr during specific phases of reach-and-grasp behavior, suggesting that pauses in pallidal firing could gate or coordinate the precise timing of movements (Falasconi et al., 2025).

However, a longstanding and ubiquitous observation is that the majority of pallidal projection neurons *increase* their firing rates with movement. Neurons with movement-related increases in firing have been reported in rodent SNr (Bryden et al., 2011; Fan et al., 2012; Gulley et al., 2002; Jin & Costa, 2010; Lintz & Felsen, 2016; Rossi et al., 2016), cat SNr (Joseph & Boussaoud, 1985; Joseph et al., 1985), bird pallidum (Goldberg & Fee, 2012), monkey SNr (Handel & Glimcher, 1999; Magariños-Ascone et al., 1992; Sato & Hikosaka, 2002; Schultz, 1986; Wichmann & Kliem, 2004), monkey GPi (Aldridge et al., 1980; Anderson & Horak, 1985; Brotchie et al., 1991; Georgopoulos et al., 1983; Mink & Thach, 1991a; Mitchell et al., 1987; Mushiake & Strick, 1995; Turner & Anderson, 1997) and even human GPi (London et al., 2026). In most reports, movement-related “increase” neurons outnumber “decrease” neurons in the SNr and GPi, often vastly so. These projection neurons are GABAergic – why would the basal ganglia increase their inhibitory output at a time when neurons throughout the motor pathways become more active? One possibility is that, simply by adding a sign change, “increase” neurons could activate downstream targets in parallel with decrease neurons. For example, increase neurons could preferentially innervate inhibitory neurons in target structures (Kaneda et al., 2008), or could secondarily excite downstream neurons through post-inhibitory rebound (Goldberg & Fee, 2012; Villalobos & Basso, 2022). In this view, increase and decrease neurons would act congruently to disinhibit their synaptic targets.

Alternatively, a longstanding hypothesis is that increase and decrease neurons act *antagonistically,* to focus or *select* actions (Alexander & Crutcher, 1990; Mink, 1996; Mink & Thach, 1993). For example, Mink and Thach (1991a) reported that 71% of movement-modulated neurons in monkey GPi increased their firing rate around the time of a visually cued wrist flexion or extension, whereas only 29% decreased. In addition, the distribution of the latencies of neural modulation (increase or decrease) largely overlapped the latencies of electromyographic (EMG) activity in agonist muscles. Given the preponderance of “increase” neurons – which presumably inhibit downstream targets – and the relatively late timing of neural modulation, the authors suggested that GPi output was unlikely to play a role in movement initiation. Rather, the decrease neurons might select the *desired* movement by disinhibition, whereas the increase neurons might inhibit *competing* movement plans (Mink, 1996; Mink & Thach, 1991a). This could provide a sharpening of movement selection, analogous to how antagonistic center-surround receptive fields produce spatially sharper responses in sensory neurons (Alexander & Crutcher, 1990; Mink & Thach, 1993). Moreover, for any desired movement, one could imagine a far larger number of potential competing movements; thus, more SNr/GPi neurons should increase their activity than decrease – as is commonly observed.

Subsequent findings in the basal ganglia are often interpreted in light of the action-selection/focusing hypothesis. For example, co-activation of direct and indirect pathway striatal spiny projection neurons (dSPNs and iSPNs) has been suggested to play a role in action selection, whereby pro-movement dSPNs could promote the desired movement and anti-movement iSPNs could suppress unwanted movements, presumably through their modulation of SNr/GPi (Barbera et al., 2016; Chen et al., 2021; Cruz et al., 2022; Cui et al., 2013; Gerfen, 2023; Isomura et al., 2013; Klaus et al., 2019; Markowitz et al., 2018).

The action-selection hypothesis is intriguing but lacks direct supporting evidence. In particular, “increase” and “decrease” neurons have only been defined functionally. It is possible that they constitute distinct neuronal subpopulations differing in terms of inputs, outputs, genetic identity, etc. However, a parsimonious view of the model is that increase and decrease neurons are two sides of the same coin: an SNr/GPi neuron that *decreases* its firing to specifically disinhibit movement *X* when it is desired could *increase* its firing to suppress movement *X* when different actions are desired (Falasconi et al., 2025; Schroll & Hamker, 2013). This *bidirectional* response pattern could be viewed as analogous to that of sensory neurons with antagonistic center-surround receptive fields, which are activated or suppressed depending on whether stimuli fall within the center or surround. To address this question, a main goal of this study is to examine the activity of SNr neurons during different types of movements: do different movements elicit responses of *opposite sign* in SNr neurons?

The timing of movement-related GPI/SNr activity also bears closer scrutiny. During arm movements in monkeys, the majority of GPi neurons change their firing rate *after* the earliest arm-muscle EMG activity (Anderson & Horak, 1985; Mink & Thach, 1991a). Moreover, selective lesion or reversible pharmacological inactivation of monkey GPi do not increase reaction times (Desmurget & Turner, 2008; Horak & Anderson, 1984; Mink & Thach, 1991b; Turner & Desmurget, 2010). For these reasons, it has been argued that the basal ganglia do not play a primary role in movement initiation. However, in those studies latencies were measured in animals moving as quickly as possible in response to a sensory cue. Reactive movements of this sort are typically less impacted in Parkinson’s disease (PD); PD patients have more difficulty with movements that are self-initiated (Glickstein & Stein, 1991; Martin, 1967; Russo et al., 2022; Sarma et al., 2012). Nonetheless, few studies have examined self-initiated movements in experimental animals. To address these questions, our lab introduced a controlled *self-timed* movement task (Lee & Assad, 2003). In the task, a sensory cue initiates a criterion timing period rather than triggering a rapid reactive movement; the animal must wait and then make the required movement *on its own*, without any further external cue. Previous findings from our lab and others suggest that striatal SPNs, motor cortical neurons and dopaminergic neurons of the substantia nigra pars compacta (SNc) are indeed active early enough to play a role in *initiating* self-timed movements – exactly the types of movements most affected in PD (Hamilos et al., 2021; Lee & Assad, 2003; Lee et al., 2006; Maimon & Assad, 2006; Yang et al., 2024). Thus, an ancillary goal of this study is to examine the activity of mouse SNr neurons during self-timed and self-initiated movements.

## Results

### Animal behavior in the self-timed reach task

Our main goal was to compare the activity of SNr neurons during two different types of self-timed movements. We started by comparing *reaching* vs. *licking*. We initially trained the first cohort of animals in the reach task alone. A second cohort was subsequently trained to perform both reach and lick tasks in interleaved blocks of trials. We shall first examine the self-timed reach task.

We trained water-restricted mice to perform the self-timed forepaw reach task while head-fixed (Figure 1a). At the start of each trial, a servomotor-mounted aluminum touch bar was swung into reaching range (800 ms travel time). With the touch bar deployed, a 100-ms start-cue (6,272 Hz tone) was played to indicate the start of the self-timing interval (Figure 1b). The task required the animal to wait at least 4 seconds after the cue and then touch the bar with its forepaw. No further cue was delivered after the initial cue; thus, the animals had to self-time the movement. If the touch occurred ≥4s after the cue (correct trial), a juice drop was delivered while the lick spout (normally retracted) was advanced via servomotor (800 ms travel time) into range of the tongue. The 800 ms delay in reward delivery was designed to discourage anticipatory licking while reaching. Extra touches made after the first touch did not yield additional reward. Both the touch bar and lick spout were retracted at trial end (10 s after cue). A touch before the 4s criterion time (incorrect trial) was not rewarded and resulted in immediate bar retraction followed by time-out until the end of trial (10 s after cue). If the animal did not touch the bar (no-move trial), it was retracted at trial end (10 s after cue). Thus, regardless of outcome, all trials ended 10s after the cue.

**Figure 1.**
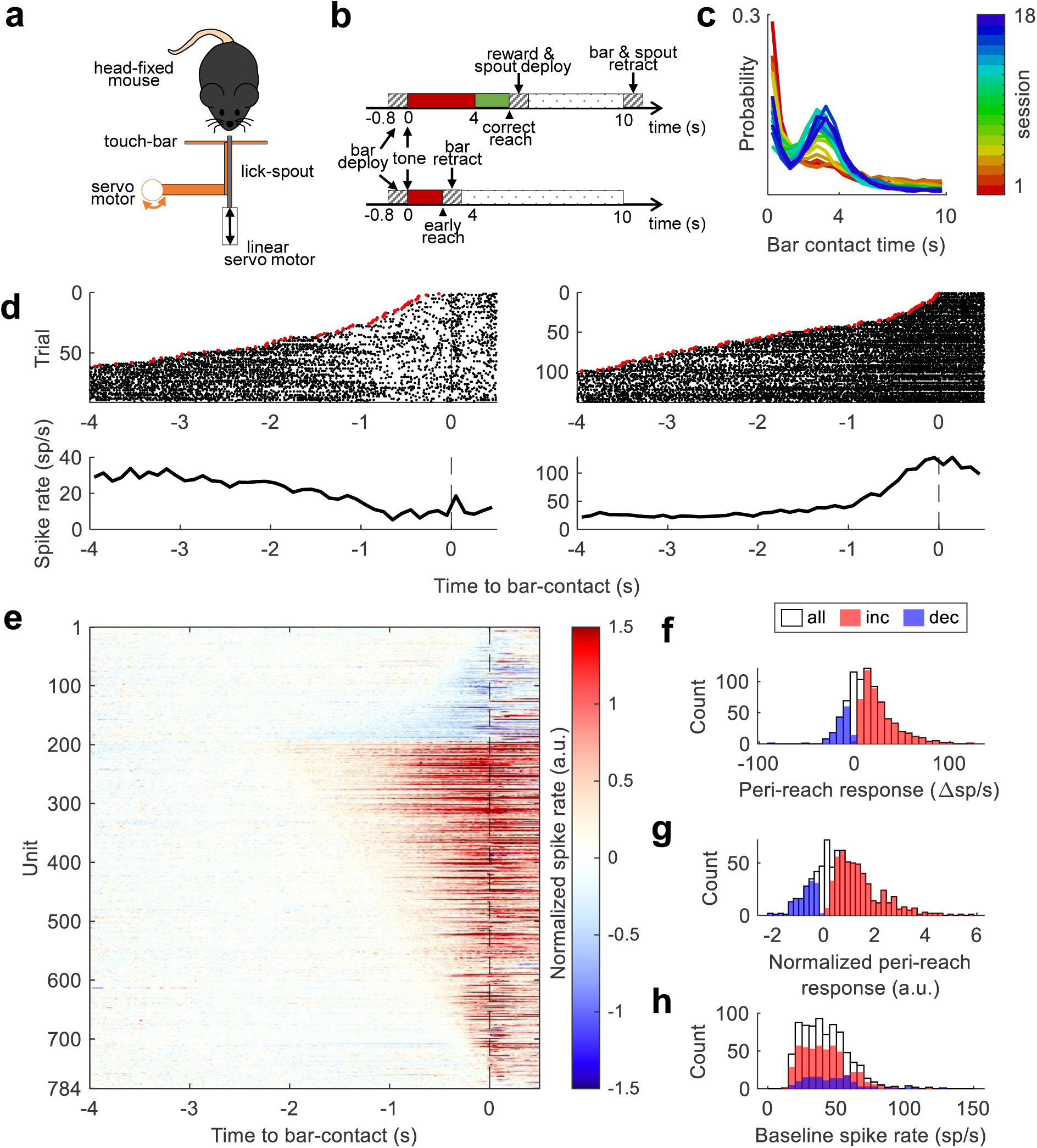
SNr neural activity in the self-timed forepaw reach task. **a.** Schematic of the self-timed forepaw reach task. **b.** Example time course of a rewarded (top) vs. unrewarded (bottom) trial, contingent on time of first reach. **c.** Distribution of movement times (i.e. first bar-contact after cue) relative to the start cue, shown in different colors for the first 18 sessions (n=29 animals). **d. Top:** spike rasters for two putative SNr neurons in the self-timed reach task, aligned to time of bar-contact; trials are sorted by movement time; spikes are not shown before the start cue (red dot); **bottom:** peri-event time histograms (PETHs) aligned to bar-contact time, for the same two SNr units. **e.** Normalized peri-movement PETHs (25ms binned spike counts) for 784 SNr units in the self-timed reach task, visualized as a heat map. Spike rates are normalized on a per-unit basis, relative to the mean and standard deviation during the “baseline window” ([−4, −2] s before bar-contact). Units are ordered by the sign and onset latency of the peri-reach response. **f.** Distribution of peri-movement response, measured as difference between the peri-movement spike rate vs. baseline spike rate ([−0.3, 0] s and [−4, −2] s relative to bar-contact, respectively). **h.** Distribution of peri-movement responses normalized to the mean and S.D. of baseline activity. **h.** Distribution of median spike rate during intertrial intervals. **f-h:** white: all SNr units (n=784); red: increase units (n=496); blue: decrease units (n=145).

In the first training session, animals were rewarded for immediately touching the bar. The reward criterion-time was gradually raised from 0.1 s to 4 s over the first 3-5 sessions. Animals learned to delay their movements as training progressed (Figure 1c). We evaluated the behavioral performance by inspecting the distribution of movement times for each session. Most trained animals generated a Gaussian-like distribution with a mode ∼4 seconds after the cue (Figure 1—Figure supplement 1). Interval estimation in animals typically shows considerable variability around the criterion time (Gallistel & Gibbon, 2000; Hamilos et al., 2021; Lee & Assad, 2003), in contrast to reactive movements, which occur with a short, stereotypical latency after the “go” cue (Lee & Assad, 2003). Thus, we focused on trials where animals moved ≥2 s after the cue, since these are unlikely to be reactive movements prompted by the cue.

### SNr activity in the self-timed reach task

We recorded single-unit extracellular activity in SNr using either chronically-implanted microwire bundles (adapted from Starkweather et al., 2017) r acutely-implanted multi-electrode silicon probes (Steinmetz et al., 2021; Yang et al., 2020). Recordings were obtained unilaterally in 29 female and male animals (17 right hemisphere, 12 left hemisphere) while they performed the self-timed reach task. To exclude dopaminergic neurons, which are also sparsely located in SNr (typically discharging at 6 sp/s; Fan et al., 2012), we selected well-isolated units with a minimum baseline firing rate of 15 sp/s as putative GABAergic SNr neurons. 784 units were considered putative SNr projection neurons (median baseline spike rate 39.7 sp/s, median absolute deviation 11.7 sp/s; the number of animals, recording sessions, and putative SNr neurons involved in the self-timed reach task, as well as subsequent experiments, are listed in Table 1).

**Table 1.** Number of SNr units and recording sessions, grouped by cohort (row) and experiment (column). ^†^ is a subset of *, as sessions with <20 self-timed lick trials were excluded.

| Cohort | Self-timed reach | Self-timed reach vs. lick | 2-target reach | 4-target reach | Spontaneous reach | Spontaneous reach vs. lick |
| --- | --- | --- | --- | --- | --- | --- |
| A (8M 3F) | 291 units<br>113 sessions |  |  |  |  |  |
| B (8M 10F) | 493 units *<br>109 sessions | 427 units †<br>76 sessions |  |  |  |  |
| C (2M 3F) |  |  | 179 units<br>10 sessions | 61 units<br>3 sessions |  |  |
| D (1M 2F) |  |  |  |  | 89 units<br>8 sessions |  |
| E (2M 4F) |  |  |  |  |  | 1,225 units<br>39 sessions |

During the post-cue/pre-movement period leading up to the forepaw reach, many SNr units either increased or decreased their firing rate. The pre- and peri-movement response time-courses of two SNr units are shown in Figure 1d, aligned to the time of touch-bar contact. In both examples, and in the majority of SNr neurons that we recorded, the average pre-movement activity changed gradually, starting hundreds of milliseconds before the movement. This pre-movement pattern of activity was reminiscent of “ramping” activity that our lab previously observed for self-timed movement tasks, in striatum (Lee & Assad, 2003; Lee et al., 2006), parietal cortex (Maimon & Assad, 2006) and SNc (Hamilos et al., 2021).

Peri-event time histograms (PETHs) of all 784 SNr units are shown as a heatmap in Figure 1e. The “baseline” spike rate of each SNr unit was estimated over the [−4 s, −2 s] window before bar contact. Because the forepaw was already moving by 200 ms before the touch-bar contact (see Figure 2), we calculated “peri-movement” activity over the [−0.3 s, 0] window before bar contact, which generally captured the peak change in activity for individual units. Using an early-shifted “pre-movement” window of [−0.5 s, −0.2 s] yielded similar results in all subsequent analyses.

**Figure 2.**
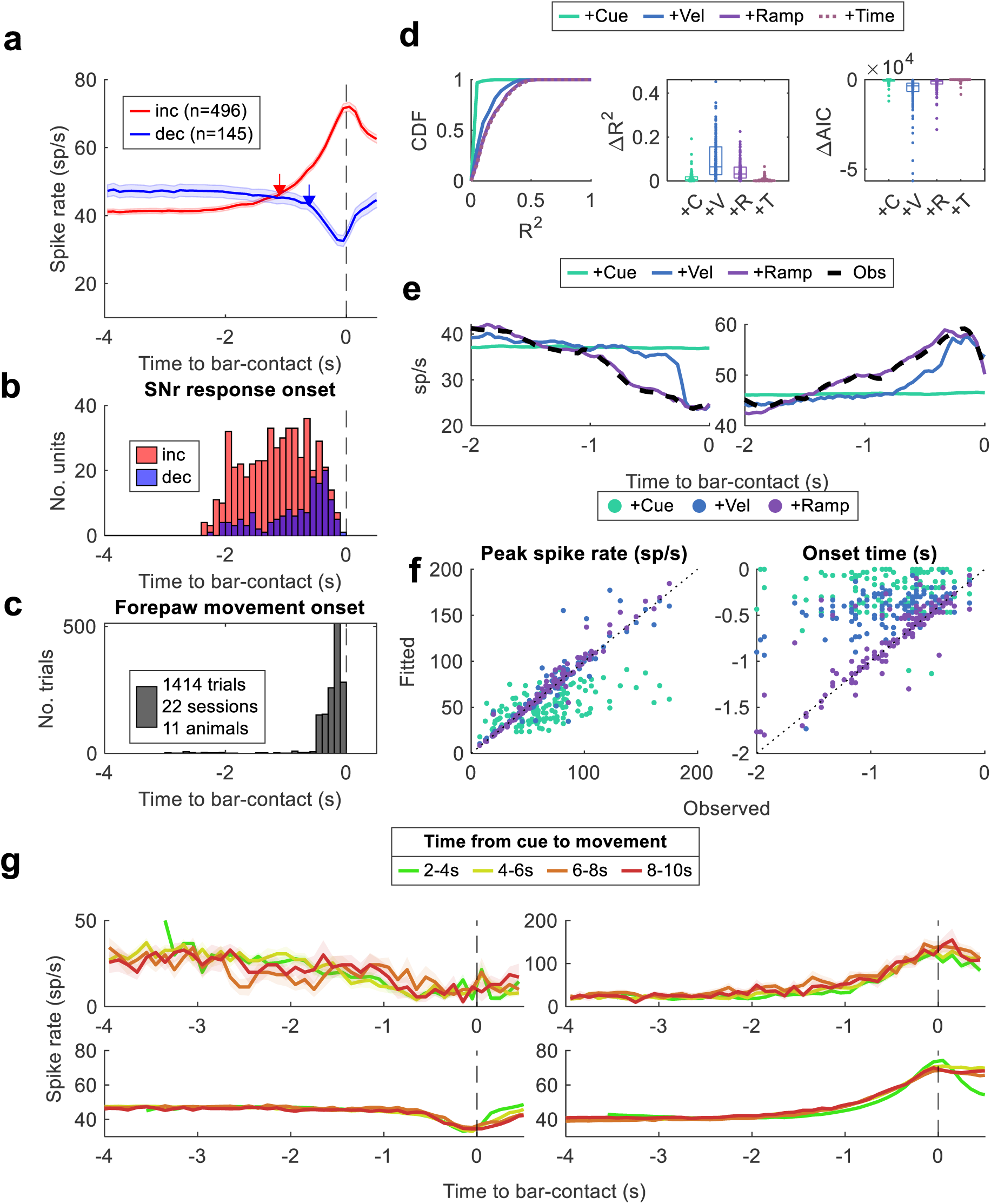
The timing of SNr neural activity in the self-timed reach task. **a.** Average PETH across increase units (red, n=496) or decrease units (blue, n=145); arrows indicate median neural response onset time (−1.10 s and −0.61 s for increase and decrease units, respectively). **b.** Distribution of neural response onset time for increase (red, n=496) and decrease (blue, n=145) units. **c.** Distribution of video-detected forepaw movement onset times relative to bar-contact. **d.** Improvement of model performance by iteratively adding predictors to the nested-GLM; **left:** cumulative distribution of explained variance (R^2^); **middle:** additional variance explained (ΔR^2^); **right:** change in Akaike information criterion (ΔAIC). Jittered dots represent individual SNr units (n=163), box plot represents the median & 25/75 percentiles. **e.** Fitted (colored traces) vs. observed (black dashes) movement-aligned PETH for two example SNr units. **f.** Fitted (y-axis) vs. observed (x-axis) PETH statistics; points represent 163 SNr units; colors represent different iterations of the nested-GLM; **left:** highest (or lowest) spike rate in the window [−0.5, 0] s before bar contact; **right:** onset latency (relative to bar-contact time) of neural response. **g.** PETH grouped by movement time; **top:** two example SNr units; **bottom:** grand average across 496 increase units (left) and 145 decrease units (right).

We measured the relative amplitude of peri-movement responses by subtracting the baseline rate (Figure 1f), or by normalizing to the mean and standard deviation of the baseline rate (Figure 1g). 641/784 (82%) SNr units showed significant pre-movement spike-rate modulation relative to baseline (permutation test, p<0.01; see Methods). Of these 641 units, 496 units (77%) increased pre-movement firing and 145 (23%) decreased pre-movement firing. The prevalence of “increase” units over “decrease” units is similar to that described in many previous studies in SNr/GPi (e.g., Mink & Thach, 1991a).

The decrease population had a significantly higher median baseline spike rate (44.6±12.1 sp/s) than the increase population (39.7±11.0 sp/s; one-sided Wilcoxon rank-sum test; p<0.0023). Nonetheless, there was a broad overlap in the distributions of baseline-firing rates (Figure 1h), and baseline rates were not a reliable basis for classifying increase vs. decrease SNr units (e.g., area under the receiver-operator curve = 0.58).

### SNr activity modulation precedes movement onset

Many SNr units showed spike-rate modulation starting hundreds of milliseconds before movement, manifesting as a gradual downwards (or upwards) “ramp” in the activity of individual SNr units, as well as in population-average activity (Figure 2a). For the 641 significantly modulated units, we quantified the time of onset of neural modulation relative to the time of touch-bar contact by determining when trial-averaged spike rates first deviated ≥0.25 SD from the mean continuously for 50 ms (see Methods). The distribution of response-onset times was broad, with many units showing modulation arising hundreds of ms before the touch-bar contact (Figure 2b). 98% of SNr units had a response onset >200 ms before touch-bar contact, and the tail of the distribution extended beyond 2000 ms. The median response-onset time was −1104 ms for increase units, and −615 ms for decrease units. However, these times were relative to touch-bar contact; the animals had to first lift their forepaws and reach before contacting the touch-bar. To determine the time of forepaw-movement initiation, we used marker-less pose estimation from video data (DeepLabCut; see Methods). Among 1,414 self-timed reach trials (11 animals, 22 recording sessions), the median movement-onset time was 175 ms before touch-bar contact, and 90% of forepaw movements were initiated within 450 ms before contact (Figure 2c). The median neural-onset times were significantly earlier than the median movement-onset time, for both increase and decrease populations (p < 8.7×10^−194^ and p < 2.4×10^−54^, respectively; one-sided Wilcoxon rank-sum test). We also measured neck-muscle EMG activity in one animal (3 sessions, 336 trials). The forepaw reach reliably evoked a signal in the neck muscles. On average (across 336 trials), EMG activity onset (when EMG activity exceeded 0.25 SD for 2 consecutive 25-ms sample points; see Methods) preceded video-detected movement-onset (when forepaw displacement exceeded 0.25 SD for 2 consecutive 25-ms sample points; see Methods) by 199 ms. Even when we artificially lengthened the (video-detected) forepaw movement onset latency by 200 ms to account for the EMG offset, the median neural-onset times still significantly preceded the median movement-onset time (p < 1.2×10^−136^ and p < 2.1×10^−18^ for increase and decrease units, respectively; one-sided Wilcoxon rank-sum test), and many neurons showed firing-rate modulations that preceded the earliest EMG activity by hundreds of milliseconds.

The early onset of firing-rate modulation relative to movement suggests that SNr neurons could play a role in initiating self-timed movements. However, SNr and other motor neurons respond to “jittery” spontaneous body movements on a much shorter timescale (Hasnain et al., 2025; Rossi et al., 2016). We indeed observed phasic SNr responses to spontaneous limb, torso, and orofacial movements during intertrial interval. Although our subjective impression was that mice tended to “settle down” at the start of self-timed movement trials, it was possible that the animals made more frequent jittery movements as trials progressed, resulting in a gradual ramping of neural activity in SNr.

To examine the possibility that SNr activity could be explained by jittery movements alone, we collected high-quality bilateral video data for 6 animals (7 sessions, 163 SNr units recorded) and used marker-less video tracking to determine the position and velocity of all four paws, the nose, and the midpoint of the spine (see Methods). When averaged across trials, the velocity of all body parts was close to zero until the initiation of the forepaw reach (Figure 2—Figure supplement 1c). This suggested that the ramping neural modulation that preceded movement initiation was likely not due to a gradual build-up of jittery body movements.

To examine this question on a trial-by-trial basis, we constructed a series of nested generalized linear models, i.e., Poisson regression (Pillow, 2007), using four categories of predictors to estimate spiking rates for individual SNr neurons. 163 SNr units were modelled independently on a unit-by-unit basis. “Cue” (C) predictors were discrete “trial-start” events convolved with 400-ms wide cosine kernels with temporal offsets ranging from −600 to 600 ms (negative offsets were included because touch-bar deployment, which was audible, began 800ms before the cue). “Velocity” (V) predictors included video-derived movement velocities of the four paws, the nose, and the midpoint of the spine, with temporal offsets ranging from −200 to 200 ms. “Ramp” (R) predictors included a series of linear ramping signals with different slopes, all terminating at the time of touch-bar contact but with different onset times ranging from 100ms to 2000ms before touch-bar contact. Finally, the “trial-progression” (T) predictor was a single linearly increasing signal beginning at cue onset and ending at touch bar contact.

Using a nested approach, we iteratively added predictors to the GLM and asked whether they improved model performance (Figure 2d), as measured by variance explained (R^2^) or Akaike information criterion (AIC). The “null” model was a constant reflecting the average spiking rate across the whole session. The “null + cue” (+C) model accounted for cue-responses in a small number of units but performed poorly overall. The “null + cue + velocity” (+V) model could predict the sign and amplitude of pre-movement modulation but failed to predict the early onset of the ramp (Figure 2f). In contrast, the “null + cue + velocity + ramp” (+R) model not only explained additional variance (Figure 2d) but also allowed for significantly better reconstructions of the pre-movement PETH (Figure 2e shows reconstructions from two example SNr units). Finally, the addition of the “trial-progress” (+T) signal did not further improve model performance (Figure 2d). Thus, we did not find evidence that a gradual ramp-up of jittery random movements could account for the early onset of spike-rate modulation in SNr neurons. Rather, the start of the ramping activity was related to the initiation of the upcoming self-timed movement in some way.

### Slope of ramping activity in SNr is timing-independent

Previous work from our lab showed upward-ramping activity among dopaminergic neurons in the SNc within a self-timed lick task. The average ramping activity began at the start-timing cue and continued until the time of movement. The slope of the ramping varied according to (and was predictive of) when animals chose to move on a trial-by-trial basis, whereby steep slopes were associated with early movements, and shallow slopes with later movements, as if the neural activity rose toward some systems-level threshold (Hamilos et al., 2021).

In contrast to our previous findings in the SNc, the onset and slope of ramping activity in SNr neurons was much more stereotyped when aligned to the start of movement (Figure 2g). To quantify this effect for each SNr unit, we grouped trials into 4 bins based on movement time (2-4, 4-6, 6-8, and 8-10 seconds) and then constructed movement-aligned PETHs for each of the four movement-time bins. For each neuron, we averaged the Euclidean distances between each pair of PETH traces (six pairwise comparisons for four bins) during the pre-movement period ([−2, 0] s prior to bar contact (to avoid reward-related differences that arose following the movement). We compared that average distance to the permuted distribution of average distances obtained from repeatedly shuffling trial identity (see Methods). Only 30/641 (5%) units showed significant differences among the timing-dependent PETHs (p < 0.01). Even for those few units, individual inspection revealed that detectable differences were most prominent at the beginning of trials, likely reflecting cue-related responses on short trials rather than pre-movement activity (Figure 2—Figure supplement 1d). Thus, we did not find evidence that ramping activity of SNr neurons varied with the timing of the movement. Rather, the ramping activity for each SNr unit generally exhibited a stereotyped slope and onset when aligned to the start of movement.

### SNr activity precede spontaneous movements

For both increase and decrease units, we found robust ramping activity (on average) arose hundreds of ms before the earliest detection of forepaw movement. This early neural modulation suggests that the SNr could, in principle, be involved in movement initiation. However, ramping activity could be an idiosyncratic feature of the timing requirement of our self-timed movement task and might not be present for “spontaneous” movements. In particular, the self-timed movement task required animals to *withhold* movement until the criterion time. Firing of inhibitory SNr neurons might play a role in active withholding, similar to how striatal iSPNs activated during a “dynamic action suppression” task (Cruz et al., 2022). At least for the increase neurons, the gradual ramping up in activity might counteract the animal’s increasing urge to move throughout the timing interval.

Several observations argue against this possibility, First, if ramping activity of increase units acts to withhold movement, we might expect that their activity should abruptly *decrease* just before movement initiation, as if withdrawing the brake. However, the SNr units in our sample generally showed monophasic responses: the sign of the peri-movement response was consistent with the sign of the ramping activity (Figure 1e; Figure 2—Figure supplement 1a). Second, at the start of each trial in the reach task, the animals consistently made an overt “flinching” movement as the servomotor began to deploy the touch-bar. This flinching often evoked a phasic response from SNr units. Invariably, the sign of this phasic cue-response matched that of the ramping activity (Figure 2—Figure supplement 1a & b): units that ramped up/down during the self-timed interval generally showed a phasic increase/decrease in activity in response to the overt flinch, respectively. These combined observations suggest that the ramping activity during the self-timed movement task was related to the upcoming movement *per se* rather than *withholding* that movement.

Notwithstanding, it is still possible that the ramping activity is a peculiar feature of self-timed movements and is not a general feature of “spontaneous” movements. To examine this question, we designed a spontaneous reach task that eliminated the explicit timing requirement (Figure 3a). Animals were allowed to move at any time, and touching the bar always resulted in a rapid “cycling” of the touch bar – immediate retraction followed by immediate redeployment – requiring a total of 1.4 s. A randomized timeout period started after any rewarded touch. During the timeout, reaches were not rewarded and resulted in immediate cycling of the touch-bar. The end of the timeout interval was not signaled to the animal. The first reach after the timeout was rewarded. The lick-tube was presented for 3-4 s for reward collection, followed by the next timeout interval. Importantly, the duration of the timeout period was randomly drawn from an exponential distribution, with a time constant of 20 s. The exponential probability was chosen because the instantaneous probability of the event of interest does not change with elapsed time (i.e., flat hazard function).

**Figure 3.**
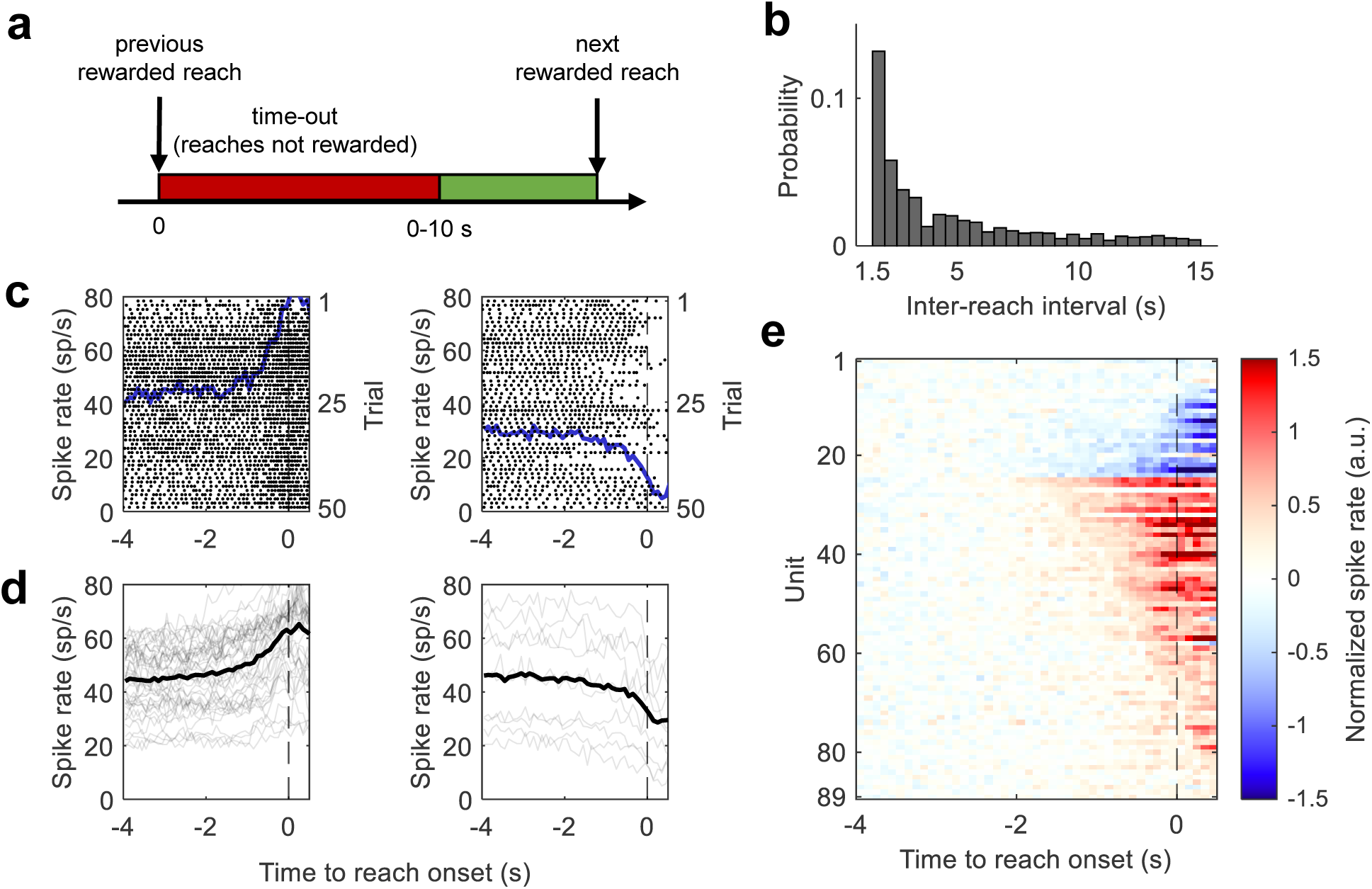
SNr neural activity in the spontaneous reach task. **a.** Time course of a spontaneous reach trial. The timeout is randomized between 0 and 10 s according to an exponential distribution. **b.** Distribution of inter-reach intervals (time between bar contacts; not shown beyond 15 s; n=1,164 trials from 3 animals, 8 sessions; individual sessions resemble the aggregate). **c.** PETH (blue trace) and spike raster (black dots) aligned to movement onset, for two example SNr units. For clarity, rasters show every 5th spike across 50 randomly selected trials. **d.** PETH of increase units (left, n=43) and decrease units (right, n=10). Dark black traces represent population average. **e.** Heat map visualization of PETHs for 89 units. Units are ordered by response sign and onset latency. **c-e:** trials with short (<6 s) inter-reach intervals are excluded.

We trained three naïve mice to perform the spontaneous reach task. The animals had never been trained in the self-timed movement task. Across all 8 sessions in the 3 animals, the distribution of inter-reach intervals declined monotonically, resembling an exponential distribution (Figure 3b) rather than the peaked distribution we observed for animals trained in the self-timed movement task. This is consistent with the animals not using a timing strategy.

We recorded from 89 putative SNr neurons in the 3 animals. The pre-movement modulation of spiking activity is shown in Figure 3c-e. A subset of 47/89 units exhibited significant pre-movement modulation relative to baseline (p<0.01, permutation test): 41 units (87%) increased in firing rate, and 6 units (13%) decreased in firing rate (Figure 3d-e). On average, the pre-movement modulation of SNr activity for spontaneous reaches resembled a gradual ramp, with most units increasing activity and a minority decreasing activity, very similar to the ramping activity we observed in the self-timed movement task (compare with Figure 1). This suggests that the pre-movement ramping is not specific to explicit timing tasks, and thus that ramping activity is unlikely to represent timing-related active withholding or suppression of movement. (An additional 6 animals were trained in the spontaneous task, but alternating between reaching and licking. Similar results were found – see below.)

### Interleaved reach vs. lick task to test the sign of neural responses in SNr

In the self-timed reach task, the majority of modulated units increased their firing rate during the pre-movement period, while only a minority of units decreased their firing. According to the action-selection hypothesis, the small contingent of decrease neurons could be selective for the forepaw movement, with their decreased firing disinhibiting the reaching movement. It follows that the larger population of increase neurons (during reaching) were selective for a *different* type of movement not involved in forepaw reaching. Their increased firing would thus have inhibited those unwanted movements during reaching. In the simplest view of this action-selection scenario (Mink & Thach, 1993), if animals were to perform an alternative movement – licking, for example – the neurons that previously decreased firing during reaching would now increase their firing, so as to inhibit forepaw movements, while a subset of the neurons that previously increased their firing during reaching – i.e., those that are selective for licking – would now decrease their firing to disinhibit licking. In this view, some individual SNr neurons should show *opposite signs* of response during the two types of movements (Schroll & Hamker, 2013).

To test this hypothesis, we trained a subset of 16 animals to perform two types of self-timed movements – forepaw reach vs. lick – in interleaved blocks of 50-75 trials within the same session. The reach task was as described above (Figure 1b), with the lick-spout remaining retracted until after a correct reach was made, to discourage anticipatory/spontaneous licking. The lick task (Figure 4a) largely resembled the reach task except for two key differences: 1) the touch-bar was retracted out of reach throughout the trial, and the lick spout was deployed (instead of the touch bar) at the start of trial; and 2) animals were rewarded if the first *lick* occurred 4-10 seconds after the start cue. In both tasks, servo motors deployed or retracted the touch-bar or lick-spout over 0.8 s. For both tasks, animals were only rewarded for the first correctly timed movement (reach or lick, depending on the task) in each trial; moving more than once did not yield more rewards. Early contact (before 4 s) with the touch bar or lick spout resulted in their immediate retraction and a timeout period. All trials (regardless of outcome) were 10 seconds in length, so there was no advantage to making early movements.

**Figure 4.**
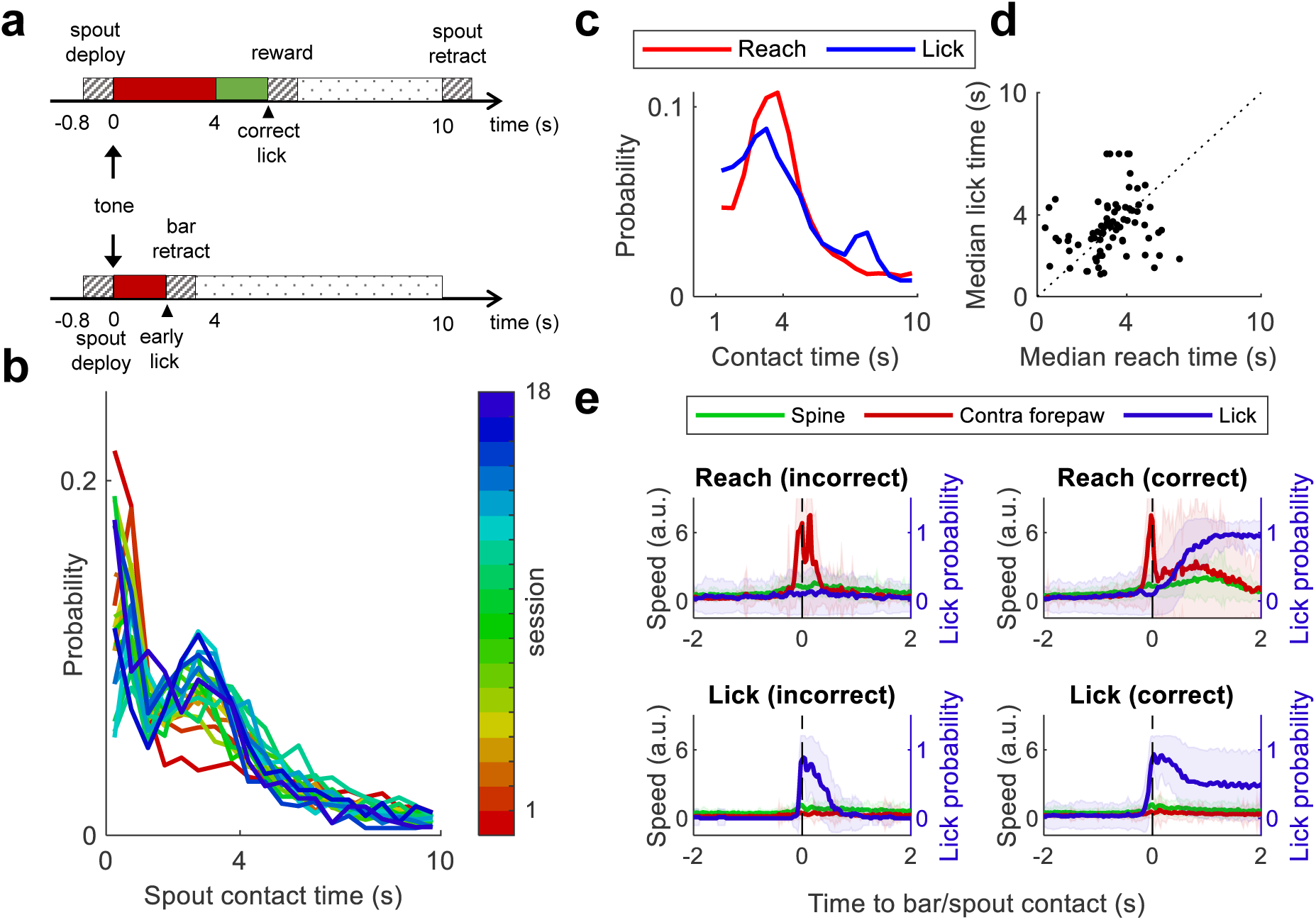
The self-timed lick vs. reach task. **a.** Example time course for a rewarded (top) or unrewarded (bottom) self-timed lick trial, contingent on lick time. For the reach task see Figure 1b. **b.** Distribution of movement times (i.e. first spout contact after cue), shown in different colors for the first 18 sessions that included self-timed lick trials (aggregate across 16 animals). **c.** Distribution of movement times across 11,238 self-timed reach trials (red) vs. 9,811 self-timed lick trials (blue). Reactive movements (made ≤1 s after cue) are not shown. **d.** Median reach time vs. median lick time relative to start cue for 79 sessions. **e.** Trial-averaged normalized movement speed of the spine (green) and the contralateral forepaw (red), as well as tongue protrusion probability (blue), derived from video analysis. Plots are shown separately for incorrect/unrewarded reach trials (n=71), correct/rewarded reach trials (n=185), incorrect/unrewarded lick trials (n=159), and correct/rewarded lick trials (n=176). Shaded area represent standard deviation.

Naïve animals were initially trained to perform only the reach (or lick) task, then learned to perform both reach and lick tasks in interleaved blocks of 50-75 trials within the same session. Within the same session, well-trained animals were required to perform at least 20 self-timed reach trials and 20 self-timed lick trials for which the movement occurred ≥2 s after the cue. Animals reached asymptotic behavior within 7-14 days of training (Figure 4b), exhibiting a broad distribution of movement times in both the reach and lick tasks (Figure 4c-d; Figure 4—Figure supplement 1).

Importantly, we designed the interleaved reach and. lick tasks to elicit an “isolated” reach or lick movement, by 1) by retracting the touch-bar out of reach throughout lick trials; and 2) delaying lick-spout deployment on reach trials. Nonetheless, animals could have made anticipatory licks before or while they reached (even though the lick tube was retracted during the reach) or fidgeted their forepaw before they licked. If the animals made *both* movements on each trial – one instructed and one spurious – this would undermine the goal of the experiment. To control for this, we collected and analyzed high resolution bilateral video recordings for a subset of 6 animals (7 total sessions). Trial-averaged time courses for movement speeds of the contralateral forepaw, the spine, as well as probability of licking/tongue protrusion (including “air licks” that failed to reach the spout), are shown in Figure 4e. During the reach task, animals generally did not engage in anticipatory licking before or during the forepaw movements: licking only occurred when the lick spout was moved into range of the tongue on correct trials (Figure 4e, top right). This delay in licking would presumably allow us to distinguish neural activity related to the earlier reach from the later lick. Conversely, animals did not make detectable forepaw movements (on average) during the lick task (Figure 4e, bottom row). In addition, video-tracking did not reveal evidence of overt bodily movements (e.g., along the spine) that might be related to postural adjustments common to both movements (Figure 4e, green trace). These behavioral measurements suggest that we were able to successfully isolate licking vs. reaching between the two tasks, which should allow us in turn to distinguish reach-related from lick-related neural activity.

### Activity during reach vs. lick task is largely congruent in individual SNr units

We recorded the activity of 427 SNr units from 16 well-trained animals. Figure 5a shows spike rasters from three example units with different response patterns: 1) decreased firing in both reach and lick trials; 2) increased firing in both reach and lick trials; 3) oppositely modulated firing in reach vs. lick trials. Across all 427 units, in both tasks, we observed more increase units than decrease units (Figure 5b). The majority of SNr units exhibited *congruent* modulation during reach and lick, either increasing or decreasing their peri-movement firing rate for *both* tasks. However, a small subset of the units showed opposite signs of modulation between the two tasks.

**Figure 5.**
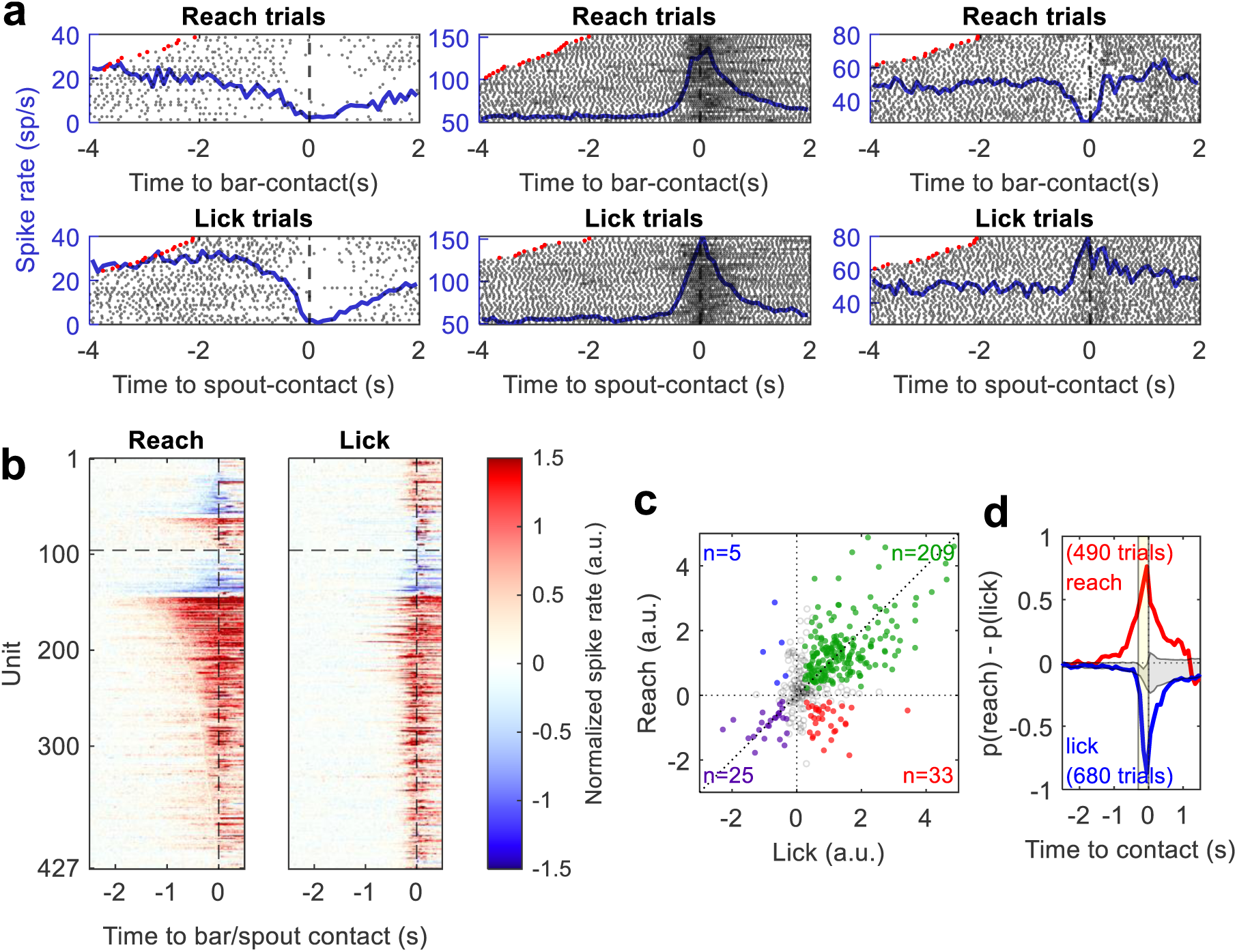
SNr neural activity in the self-timed lick vs. reach task. **a.** Raster plots of spiking activity for three example SNr units during 40 self-timed reach trials (top) vs. 40 self-timed lick trials (bottom). Trials are sorted by movement time, with red dots indicating the start cue. Black dots represent spikes (every 4th spike is shown for clarity). Blue traces represents PETHs. **b.** Heat map visualization of peri-reach (left) and peri-lick (right) PETHs for 427 SNr units arranged in the same order. The horizontal dashed line separates oppositely-modulated units (above) from congruent units (below). PETHs are normalized to baseline spike rates [−4, −2] s before movement. **c.** Normalized peri-movement activity ([−0.3s, 0] before bar/spout contact) during reach vs. lick: colored dots represent 272 units significantly modulated for both reach and lick, split into 4 subpopulations based on the sign of peri-reach vs. peri-lick responses; grey circles represent the remaining 155 units. **d.** LDA decoding of SNr population activity: decoded probability of reach subtracted by probability of lick is averaged among true reach trials (red) vs. true lick trials (blue). Yellow shaded area represents peri-movement training window: [−0.3, 0] s. Model is trained on SNr population activity from 490 reach trials and 680 lick trials (10 animals, 13 sessions, ≥10 SNr units per session, 293 total units). Grey shaded area represent 99% CI of null models trained on shuffled trials (10,000 permutations).

To quantify these findings, we averaged the peri-movement response of each SNr unit over the window [−300, 0] ms before target contact, for both tasks, and we determined if the response was statistically significantly different from the baseline activity, as described above (permutation test, p<0.01). For the two tasks separately, we classified SNr units as “increase”, “decrease”, or “unmodulated”, based on their sign of (or lack of) response. Figure 5c shows the average peri-reach vs. peri-lick responses for all 427 units. 272/427 units were significantly modulated in both tasks, allowing us to classify the sign of the response for both reach and lick (the remaining 155 units were significantly modulated in only one or neither of the two tasks). Of these, the vast majority (234/272, 86%) had congruent signs of neural modulation for both forepaw and licking movements, either increasing or decreasing firing for both tasks. Only a small minority of SNr units exhibited a sign reversal across the two tasks – 14% of units significantly modulated in both tasks; 9% of all recorded units. Across the entire population of 427 SNr units, there was a significant positive correlation between peri-reach and peri-lick modulation among all 427 SNr units (R^2^=0.29, p < 4.3×10^−34^ under null hypothesis of zero-slope; linear regression), consistent with the majority of units having a congruent sign of modulation between the two tasks.

Of special interest are the subset of SNr units that showed decreased firing during the peri-movement period. Under the simple action-selection hypothesis, the decreased firing in these units could act to disinhibit a desired movement – but then those same units should increase firing to suppress that movement when it is becomes undesired. However, even among the 93 units that showed significant *decreases* in firing in either task, only a minority showed increased firing in the other task (38/93, 41%). In conclusion, few SNr neurons showed opposite signs of firing-rate modulation between the reach and lick tasks. This finding is difficult to square with a simple model whereby SNr neurons select or de-select actions.

### Congruent SNr activity in the reach vs. lick task is not due to movement planning

Whereas most SNr units showed congruent signs of activity in the reach vs. lick task, with either increased or decreased firing in both tasks, we were initially concerned that spurious forepaw movements (during the lick task) or anticipatory licking (during the reach task) could artifactually present as congruent neural modulation. Using video-tracking, we detected neither spurious forepaw movements during the lick task nor anticipatory licking during the reach task (Fig. 4e), allowing us – at least in principle – to distinguish neural activity related to reaching and licking. However, even if *overt* movements were dissociated in time, animals could in principle still *plan* to perform both actions, and this could potentially explain the apparent congruent neural responses we observed between the two tasks. This was less of a concern for the lick task, because the touch bar was retracted out of reach of the animals throughout the entire block of lick trials: presumably, the animals did not plan to reach if there was nothing to reach to. The interleaved reach task, however, could be more confounded, because the animals could anticipate licking following the reach on correct trials. In this scenario, if a unit were selective only for licking, that unit might then show congruent pre-movement modulation between the lick and reach task, because the animal planned to lick in both cases; this could lead us to mis-characterize that unit as non-selective between the tasks. To mitigate this concern, we delayed the licking at the end of reach trials by slowly deploying the lick-tube (over 800 ms) into range of the tongue following a correct reach. We reasoned that by delaying the lick-tube deployment, potential lick-related pre-movement modulation would likewise be delayed for lick-selective units. Indeed, video tracking revealed that tongue movements were delayed in the reach task compared to the lick task (Figure 4e, top vs. bottom row). However, the pre-movement neural response was *not* systematically delayed in the reach task; if anything, the onset of pre-movement neural modulation was noticeably *earlier* in the reach task than the lick task (Figure 5b). This suggests that the pre-movement neural modulation during the reach task was indeed related to the forearm reach, not planning the upcoming lick.

### Amplitude of neural modulation in the lick vs. reach task

While the sign of neural modulation was congruent between lick and reach for most SNr units, we also examined the relative amplitude of peri-reach vs. peri-lick modulation. 188/427 (44%) units exhibited significantly different modulation amplitudes between reach and lick tasks (p<0.01; permutation test, see Methods), even though the signs of the responses were largely congruent between the two tasks. At the population level, these differences were substantive enough that we could use SNr responses to classify individual trials as reach or lick trials. We trained a linear-discriminant decoding model (see Methods) to predict the underlying trial type using population neural activity (≥10 SNr units/session, 13 sessions, 293 total units; 490 reach trials vs. 680 lick trials).

Although the models were trained on neural activity from the peri-movement period ([−0.3, 0] s prior to touch-bar or lick-spout contact), they were able to successfully decode the true movement type hundreds of ms before and after the movement (Figure 5d). Therefore, although individual SNr neurons mainly exhibited congruent signs of modulation for reach vs. lick, the population of SNr neurons still carried information about the type of movement on a trial-by-trial basis.

### Congruent SNr modulation across multiple phases of movement

In our self-timed reach vs. lick task, the vast majority of neurons increased firing during both operant movements. However, these neurons could have specific decreases in firing for other movements/motor programs that we did not test explicitly. Apropos, a recent study with mice performing a grasping task (Falasconi et al., 2025) reported that a majority of SNr neurons decreased firing during specific phases of the movement (reaching, grasping, retraction, and food consumption). We thus examined the activity of SNr neurons across additional phases of movement in our self-timed reach vs. lick task. We divided our reach vs. lick task into 5 phases: <u>reach</u> – the initial reach/arm-extension in reach trials; <u>lick</u> – the initial lick in self-timed lick trials; <u>first lick</u> – the start of a lick bout in rewarded reach trials, which typically occurred ∼1s after reaching; <u>last lick</u> – the end of a lick-bout in rewarded trials, when animals finished collecting the reward; <u>bar release</u> – arm retraction at the end of rewarded reach trials, always 10 s after the start cue when the bar was retracted. Due to electrical artifacts caused by touch sensors and servo motors for some neurons, we restricted our analyses to a subset of 298 SNr units where such artifacts were absent or could be filtered out (see Methods).

The spiking activity of two example SNr neurons are shown in Figure 6a; all 298 neurons are shown as a heat map in Figure 6b. For each of the five phases of movement, the majority of significantly modulated neurons (p<0.01, permutation test; see Methods) *increased* rather than *decreased* their firing rates, detailed as follows: reach (n=226 units, 86%/14% increase/decrease), lick (n=252, 92%/8%), first lick (n=236, 88%/12%), last lick (n=225, 90%/10%), bar release (n=238, 91%/9%). Many of these modulated SNr neurons (n=133, 45% of total) still exhibited congruent *increases* in firing rates across *all five phases* of movement. 6 neurons (2%) decreased firing rates for all 5 phases of movement, while an additional 21 (7%) decreased firing rates for two or more (but not all) phases. Notably, only 37 SNr neurons (12%) decreased firing rates for one specific phase of movement and increased for at least one other movement; an even smaller contingent of 12 neurons (4%) decreased for one phase and increased firing rates for all other phases. Thus, while some neurons showed specific decreases for one particular phase of movement, this was a relatively uncommon motif in our data set; congruent firing patterns were far more prevalent.

**Figure 6.**
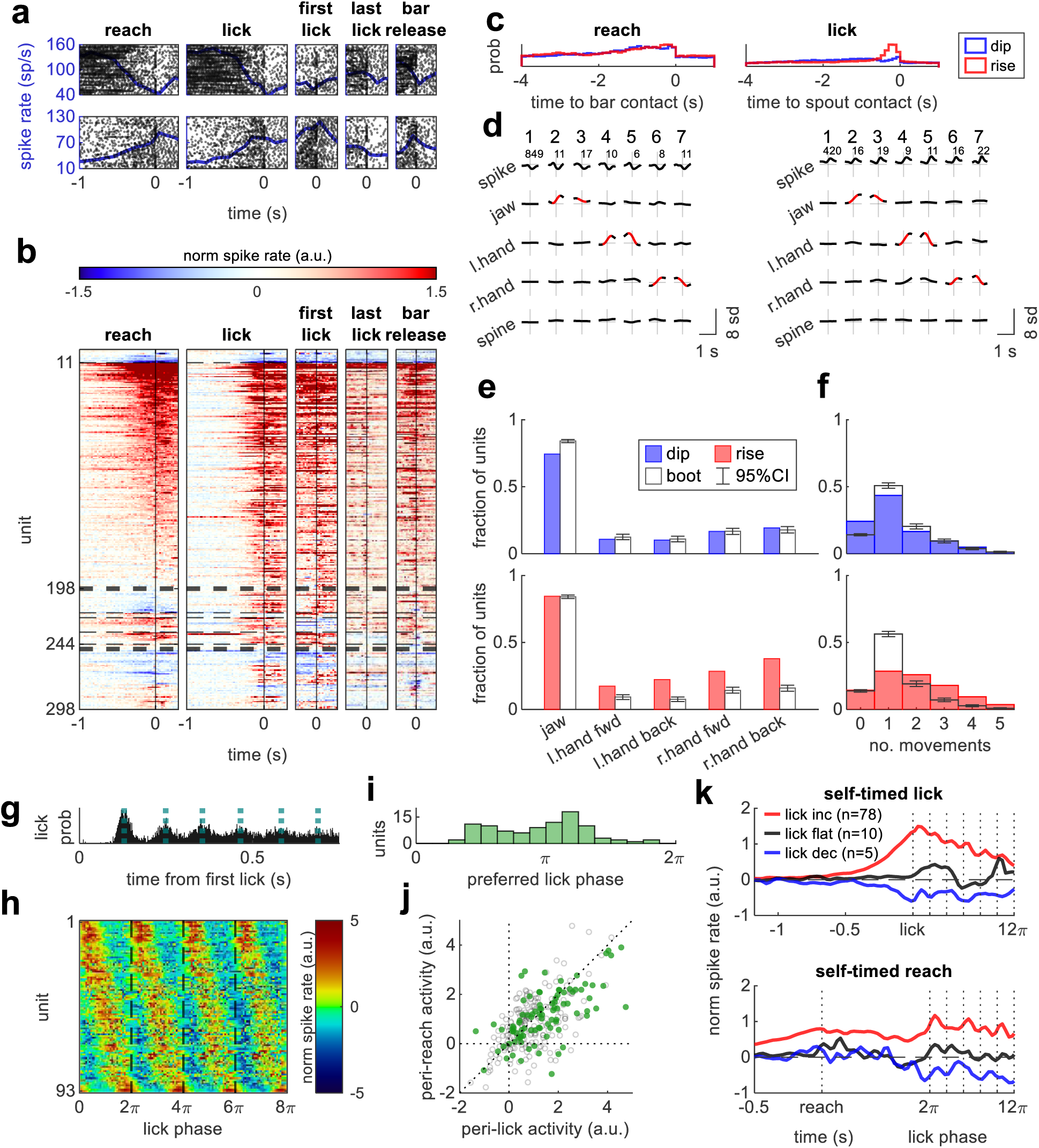
Dips in SNr firing do not encode single movements. **a.** Two example SNr units and their spiking activity (PETH in blue) aligned to 5 phases of movement: **reach**: the first bar-contact in a reach trial; **lick**: the first spout-contact in a lick trial; **first lick**: the first spout contact in a rewarded reach trial; **last lick**: the last spout-contact in a rewarded reach or lick trial; **bar release**: when the paw releases the bar at the end of a rewarded reach trial. **b.** Normalized peri-movement modulation of 298 SNr units aligned to 5 phases of movement. Units are arranged in 3 blocks, from top to bottom: congruent modulation; decrease for only one phase; decrease for 2 or more phases. White represents baseline firing [−4, −2] s before reach. **c.** the probability of transient (200-600ms) dips/rises in SNr firing rate during spontaneous reach (n=580) and lick trials (n=913). **d.** Two example SNr units whose transient dips (left) and rises (right) were clustered based on the accompanying movement profile: 1: no move, 2: jaw open, 3: jaw close, 4: left hand forward, 5: left hand back, 6: right hand forward, 7: right hand back; the number of dips/rises are noted below the cluster index; rows from top to bottom are: spike rate, jaw DV pos, left hand AP pos, right hand AP pos, spine DV pos. **e.** Fraction of SNr units (1,225 total) with >10 dips (top/blue) or rises (bottom/red) observed along with movements of the jaw (open/close combined), left hand (forward & back), or right hand (forward & back). **f.** Fraction of SNr units grouped by how movement types were observed during dips (top/blue) and rises (bottom/red). Hollow bars & error bars represent the mean & 95% CI of the bootstrapped distribution. **g.** Lick histogram aligned to the first lick in each trial (n=33,337 trials); dashed lines indicate estimated peaks in lick probability. **h.** Normalized PETHs aligned to lick-phase for 93 lick-entrained SNr units, ordered by preferred phase (0, 2*π,* etc. denote spout-contact). **i.** Distribution of phase preference among lick-entrained SNr units. **j.** Normalized peri-reach vs. peri-lick responses ([−0.3, 0] s before self-timed bar/spout contact) for 298 SNr units, of which 93 lick-entrained units are colored green. **k.** Normalized spike rates of example SNr units during rewarded self-timed lick (top) or reach (bottom); red/black/blue traces represent population average among lick-increase, lick-flat, lick-decrease SNr units, respectively. Spike rates are normalized to baseline firing [−4, −2] s before reach. Spikes after the first lick are binned relative to the licking cycle.

### Brief dips and rises in SNr firing rate are not related to stereotyped movements

Even though we found largely congruent responses throughout the entire complex sequence of movements making up the behavioral tasks, it is possible that we still did not test the “right” movement to evoke specific dips in firing. We therefore took the reverse approach: rather than looking for dips in firing with respect to particular experiment-defined movements, we detected *any* transient change in firing – dips or peaks – during and between the experimental trials, and then asked if those firing transients were, on average, associated with specific movements.

To this end, we trained an additional 6 animals in the spontaneous reach vs. lick task, but with high-resolution imaging of the entire body to more broadly detect movements. We recorded an additional 1,225 putative SNr neurons across 39 sessions. The overall proportion of experimental response patterns were very similar to that found in the previous experiments (Figure 5—Figure supplement 1). But in addition, for each SNr unit, we detected brief (200-600 ms) dips (n=683±257 per unit) or rises (n=635±211 per unit) in firing rate throughout the recording session (duration=80±14 min). These dips and rises could occur any time throughout the trial (Figure 6c) or during the ITI. For each SNr unit, we extracted dip/rise-aligned kinematics of the jaw, left hand, right hand, and spine. Dips/rises were then clustered in an unsupervised manner into 7 classes based on the accompanying movement profile (Figure 6d; see Methods). These clusters represented 7 types of movement profiles (1-“no movement on average“, 2-“jaw open”, 3-“jaw close”, 4-“left hand forward”, 5-“left hand back”, 6-“right hand forward”, 7-“right hand back”). Cluster 1 contained the vast majority of dips (89%) and rises (81%), where the position of all 4 body parts did not change on a trial-average basis, suggesting that brief fluctuations in firing rate in individual SNr neurons were unlikely to accompany specific movement patterns. Nonetheless, we analyzed more closely clusters 2-7, which captured individual direction-specific movements of the jaw and forelimbs, to potentially reveal SNr neurons selective for specific movements. To ensure the quality of the unsupervised clustering for each unit, we only included clusters for which a significant movement was observed in one and only one body part (see Methods). We also combined clusters 2&3 (“jaw open” and “jaw close”), because 30-fps video sampling was insufficient to accurately delineate the rapid direction-reversal of jaw movements.

Under the simplified action-selection model, we hypothesized that any individual SNr neuron should only dip for one specific movement profile out of the five we analyzed. We estimated the chance level with a bootstrap analysis, using randomly selected intervals throughout the session instead of true dips or rises in neural activity (see Methods). Compared to the bootstrapped null distribution, we found that the fraction of SNr units that dipped for each of the five specific movements was no higher than that expected from chance (Figure 6e, top). Furthermore, the observed fraction of units that dipped for exactly *one* movement (44%) was significantly lower than that expected from chance (below the 95% confidence interval of [49%, 53%]; Figure 6f top).

Conversely, rises in firing rate accompanying the five movement types were more commonly observed among a larger fraction of SNr units (Figure 6e, bottom). However, most of the rises were not movement-selective: 29% of SNr units showed rises for only one movement, whereas 57% of SNr units had rises for 2 or more movements (higher than chance: [28%, 32%]; Figure 6f, bottom). These combined data suggest that transient dips (or rises) in the firing rate of individual SNr neurons were generally not tied to any single stereotyped movement that we could detect.

### Lick-entrained activity modulation is common among SNr neurons

On rewarded trials, the lick spout was presented for 2-6 seconds (depending on movement time). During this period, animals performed bouts of rhythmic licking with a stereotypical frequency of 8-10 Hz (Figure 6g). We identified a sub-population of 93/298 (31%) SNr neurons whose firing was strongly entrained to the periodic protrusion/retraction of the tongue observed during lick bouts (Figure 6h; p < 0.01, permutation test on circular statistics of spiking rates; see Methods). The phase of the entrainment varied considerably among the 93 units (Figure 6i), but were broadly biased toward anti-phasic (i.e., firing less during spout contact/maximal tongue protrusion), consistent with a previous study (Rossi et al., 2016).

The periodic firing pattern of these 93 lick-entrained neurons is presumably related to licking *per se*, rather than, for example, some gross postural adjustment. We hypothesized that the subpopulation of lick-entrained neurons would also show strong preference for licking over reaching in the self-timed movement task. However, even for this subset of lick-entrained units, the signs of the peri-movement modulation between the lick and reach tasks were overwhelmingly congruent (linear regression; R^2^=0.46, p < 6.2×10^−14^; linear regression; Figure 6j). On average, the lick-entrained pattern resembled small fluctuations in spike rates superimposed on the slower, higher-amplitude modulation observed during self-timed movements, for both increase and decrease units (Figure 6k). Regardless, the observation that even lick-entrained SNr units also showed largely congruent neural responses for reach *and* lick is difficult to square with the simple action-selection model (see Discussion).

### SNr responses do not change sign for different reach directions

We observed relatively few SNr neurons with opposite modulation for reach vs. lick; however, it is possible that a putative antagonistic action-selection mechanism in the SNr would only apply to movements that are *mutually exclusive*. Arm and orofacial movements are presumably not mutually exclusive (in the sense that “walking and chewing gum” are not mutually exclusive). But *different* movements of the *same* effector are, by definition, mutually exclusive −- they cannot be performed simultaneously. We thus designed a bi-directional movement task in which animals reached towards two different targets on interleaved trials. Animals were trained to use their contralateral forepaw to touch a small metal cube (5 mm wide by 10 mm high). Upon trial start, the target was presented at one of two possible locations, either lateral or medial to the contralateral forepaw. The target position was adjusted using a linear actuator before being deployed into range using a servomotor. Animals used their contralateral forepaw for most reaches; occasional trials with reaches of the ipsilateral paw (or both paws) were excluded from further analysis.

Animals were trained to make self-initiated reaches in the spontaneous movement task described in Figure 3. The target position was changed after every 15 rewarded trials. We recorded the forepaw movements with three cameras (front/left/right) and manually traced and reconstructed the movement trajectories of both forepaws. The average reach trajectories of the contralateral forepaw indeed differed depending on target position, while the ipsilateral paw remained stationary (Figure 7a).

**Figure 7.**
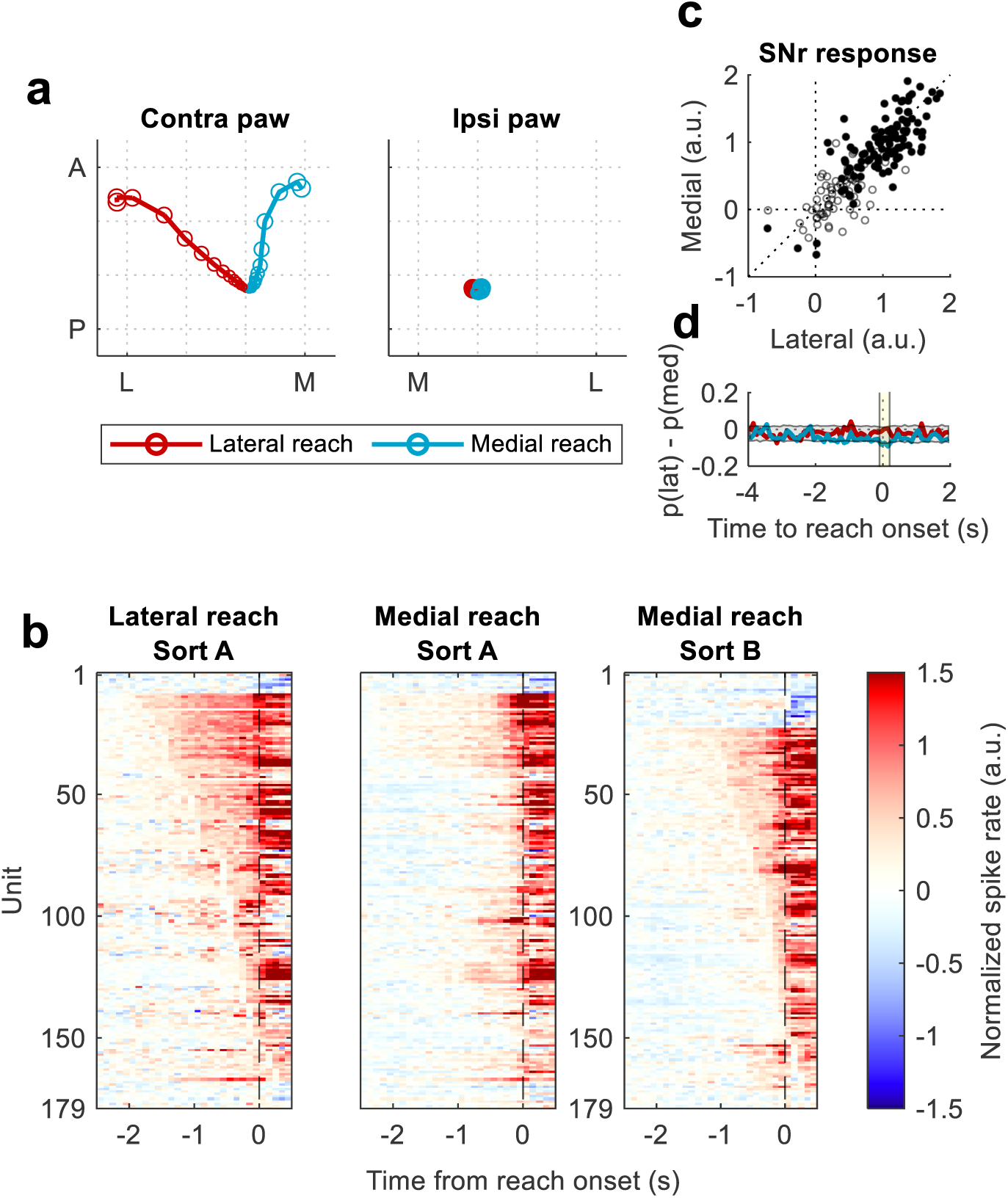
SNr neural activity in the directional reach task. **a.** Trial-averaged movement trajectories of the contralateral paw (left plot) and ipsilateral paw (right plot). Trajectories are centered relative to the resting position of the paw (i.e. 500ms prior to touch). Colors indicate whether the reach target was presented at the lateral (red, n=413 trials) or medial (blue, n=544 trials) position (3 animals, 10 sessions). Trials where animals reached with the ipsilateral paw or both paws (n=384 trials) were not included. **b.** Heat map visualization of normalized PETH for 179 SNr units during lateral reaches (left plot) vs. medial reaches (middle/right plots). Left two plots (sort A): units are ordered according to response sign/latency during lateral reach. Right plot (sort B): units are ordered according to response sign/latency during medial reach. **c.** Normalized spike rates around the time of reach onset (−100ms to 200ms) for 179 SNr units during lateral reaches vs. medial reaches, filled in dots (95/179) represent units with significant peri-movement responses for one or both reach directions (permutation test; p<0.01). **d.** LDA decoding of SNr population activity: decoded probability of lateral reach subtracted by probability of medial reach is shown for true lateral reach trials (red) vs. true medial reach trials (blue). Yellow shaded area [−0.1, 0.2] s represents training window. Model is trained on SNr population activity from 351 lateral reach trials vs. 424 medial reach trials (3 animals, 10 sessions, ≥10 SNr units per session, 165 total units). Grey shaded area represents 99% CI of null models trained on shuffled trials (10,000 permutations).

We recorded 179 SNr units (10 sessions) in 3 animals trained to perform the bi-directional reach task. The onset of arm movement was determined through video tracking. Reaches that followed a previous reach by <4s were excluded to focus on self-initiated rather than repetitive movements. For each SNr unit, we calculated the normalized trial-averaged peri-movement activity time course during lateral reaches vs. medial reaches (Figure 7b). For visualization and comparison, spike rates were normalized to the mean and SD during the [−4 s, −2 s] baseline window prior to reach onset.

Similar to the single-target reach tasks described above, we observed early neural modulation, as well as a majority of units that increased rather than decreased their pre-movement activity (Figure 7b). However, peri-movement neural modulation in SNr (within the window [−100 ms, 200 ms] relative to reach onset) was overwhelmingly similar between the two reaching targets. 95/179 (53%) SNr units exhibited significant (p<0.01; permutation test) peri-movement modulation for one or both reach directions, but none of them (0/95) exhibited opposite *signs* of modulation (Figure 7c). Furthermore, only 10/179 (6%) exhibited a significant difference in modulation *amplitude* (p<0.01; permutation test) between lateral vs. medial reaches.

We then asked whether activity in SNr could be used to decode lateral vs. medial reaches made with the contralateral forepaw. We performed linear discriminant analysis by training decoding models to predict whether the animal was making a lateral or medial reach, using population neural activity (7 sessions with ≥10 units/session, 165 total SNr units; 351 lateral vs. 424 medial reach trials). The trained models were not able to distinguish lateral vs. medial reaches: decoding performance was not significantly better than chance (p>0.01; comparing to null models trained after shuffling the trial labels; 10,000 permutations; Figure 7d). Similar results were found for a 4-direction reach task, with 3 contralateral targets and 1 ipsilateral target, with one noticeable difference: SNr neurons exhibited slightly higher firing rates (and thus presumably produced more inhibitory output,) when reaching with the ipsilateral forepaw, presumably a manifestation of the contralateral bias of forepaw motor control (Figure 7—Figure supplement 1).

In conclusion, we found no evidence that SNr neurons changed sign or amplitude of modulation for reaches made with the same forepaw towards different directions, despite the mutual exclusivity of those actions.

### Properties of SNr neurons projecting to specific motor sub-zones in superior colliculus

The superior colliculus (SC) is a major projection target of the SNr. Neurons in SNr could potentially control the selection of actions by differentially modulating SC neurons that generate specific actions. The motor SC contains sub-zones that are selective for certain types of movements (Benavidez et al., 2021; Wheatcroft et al., 2022). Circuit tracing (Lee et al., 2020) has revealed an orofacial pathway (VLS→SNr→lateral SC) separate from a skeletomotor pathway (DLS→SNr→intermediate SC). Preliminary data suggested that artificial activation of central SC could trigger forelimb movements (T. Wheatcroft, personal communication, 2026). Conversely, optogenetic inhibition of lateral SC was shown to prevent licking (Sayed et al., 2026).

We first verified the segregation of orofacial and arm movements in motor SC using optogenetic stimulation. We used two small injections of AAVs, one to express CoChR (blue-light sensitive) in central SC and the other to express ChrimsonR (red-light sensitive) in lateral SC (Figure 8a, Figure 8—Figure supplement 1a&b; also see Methods). Unilateral activation of lateral SC with red light (635 nm) triggered only jaw movements, but unilateral activation of central SC with blue light (470 nm) triggered movements of the contralateral forearm and the jaw (Figure 8b, n=3 animals, 4 sessions, 160 trials). This suggests that the representation of forearm movements is restricted to central SC, whereas the orofacial zone spans lateral and central SC. However, there are alternative explanations. First, injection of CoChR into central SC could have spread into lateral SC. However, *post hoc* histology revealed clear separation between the injection sites (Fig. S8b). Second, because ChrimsonR can also be activated by 470-nm light (albeit at a much lower efficiency than 635-nm light), 470-nm illumination could have simultaneously activated both CoChR (in central SC) and ChrimsonR (in lateral SC). This scenario is unlikely, however, because the power-threshold to trigger jaw movements with blue light (0.5 mW) was much lower than that of red light (16 mW), the opposite of what would be expected for ChrimsonR. In addition, we repeated the experiment with the injection sites *reversed* – ChrimsonR in central/forearm SC and CoChR in lateral/jaw SC (n=2 animals; see Figure 8—Figure supplement 1c). In the reverse-injection cohort, blue light only triggered jaw movements but not arm movements, even at high power (16 mW), whereas the same level of red light (16 mW) was sufficient to trigger both jaw and forearm movements (Figure 8—Figure supplement 1d). These findings argue against spurious ChrimsonR activation by blue light, and instead suggest that the jaw representation in motor SC indeed extends to both central and lateral SC.

**Figure 8.**
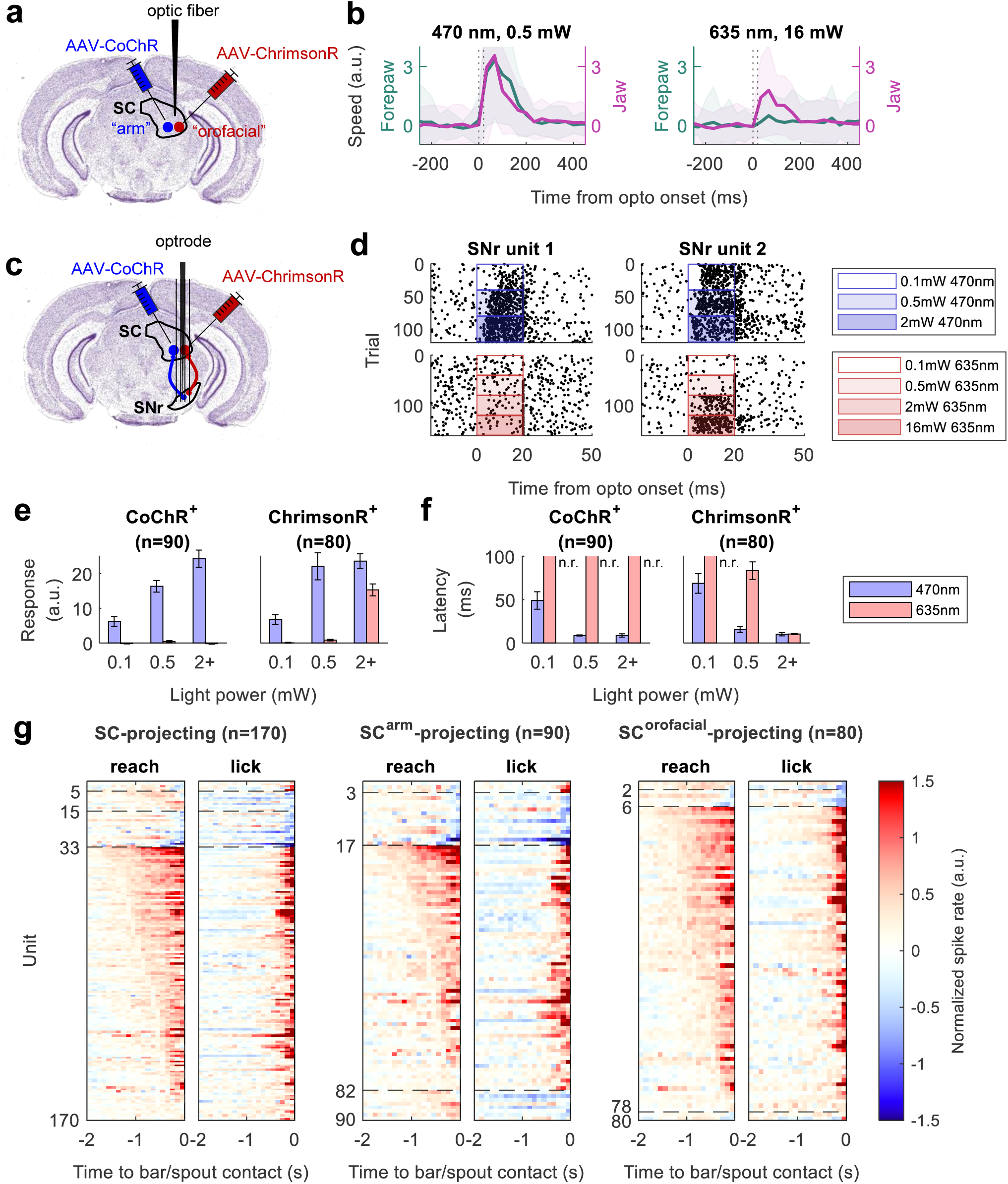
Characterization of SC-projecting SNr neurons. **a.** Diagram of two-colored SC optogenetic stimulation (adapted from the Allen Mouse Brain Atlas, 2011). **b.** Forepaw and jaw movements evoked by optogenetic stimulation of medial vs. lateral SC (averaged across 160 trials, 4 sessions, 3 animals). Das hed vertical lines indicate 20 ms laser pulse. Evoked movement speeds are normalized to baseline 0 – 250 ms before laser onset. Shaded area represents standard deviation across trials. **c.** Diagram of dual-colored optotagging of SNr neurons projecting to either the arm or orofacial zone of SC. **d.** Spike rasters of two example SNr units during 470nm and 635nm laser stimulation. Rectangles indicate the duration and power of laser pulses. **e.** Average z-scored stimulation response to 470nm vs. 635nm light (10-20ms pulses) with increasing light power; left: 90 SNr units responsive to 470nm but not 635nm stimulation; right: 80 SNr units responsive to 635nm stimulation; error bars represent 0.1SD. **f.** Average latency of stimulation response (see Methods) to 470nm vs. 635nm stimulation; n.r. – not responsive to stimulation, latency not applicable. **g.** Heatmap representation of SNr modulation during the reach task vs. lick task, shown separately for three populations, from left to right:1) putative SC-projecting SNr neurons, i.e., opto-tagged by either opsin; 2) putative arm-SC-projecting SNr neurons, i.e. CoChR^+^; 3) putative orofacial-SC-projecting SNr neurons, i.e. ChrimsonR^+^.

Based on the verified movement selectivity of lateral vs. central SC – and on the assumption that dips in firing of SNr neurons trigger specific movements – we hypothesized that 1) SNr neurons projecting to lateral (orofacial) SC should decrease firing during the lick task; and 2) SNr neurons projecting to central (forearm/orofacial) SC should decrease firing during both reach and lick tasks.

In the same cohort of animals (n=3), we looked for putative SC-projecting SNr neurons expressing either CoChR or ChrimsonR via retrograde transport of AAV vectors injected into central or lateral SC, respectively. We acutely implanted an optrode in SNr (Figure 8c) to identify SC-projecting SNr neurons via opto-tagging during the intertrial intervals of the spontaneous reach vs. lick task. We identified 170 SC-projecting SNr neurons (Figure 8d). 90 of these neurons responded only to blue light (CoChR^+^), and thus putatively projected to central SC. The remaining 80 neurons responded to both red and blue light (ChrimsonR^+^), and thus putatively projected to lateral SC. With higher light powers, the amplitude of optical responses increased (Figure 8e), while response latency decreased (Figure 8f).

We categorized the movement-related responses of the 170 SC-projecting SNr neurons during the reach vs. lick task (Figure 8g). Overall, an overwhelming majority of neurons (137/170) *increased* firing during both reach and lick tasks (Figure 8g, left panel). This pattern held when we divided the SNr neurons according to whether they projected to either central (arm/orofacial) SC (CoChR^+^) or lateral (orofacial) SC (ChrimsonR^+^): only a small minority of CoChR^+^ neurons decreased firing for either reaching (19/90) or licking (22/90) (Figure 8g, middle panel), and only a small minority of ChrimsonR^+^ neurons decreased firing for licking (6/80) (Figure 8g, right).Thus, we found scant evidence for motor-selective decreases in SNr firing that might trigger desired movements by disinhibition of specific SC motor targets. On the contrary, the vast majority of SNr neurons projecting to a movement-specific zone in SC paradoxically *increased* their firing at the same time that particular movement was executed.

### SNr functional somatotopy is not clearly defined

We found few neurons with opposite response signs for different movement types in SNr, but there may be some order in the response types if we consider the anatomical location of individual neurons within the SNr. The SNr receives convergent input from striatum: anatomical viral tracing studies (Foster et al., 2021; Hintiryan et al., 2016) have revealed parallel, spatially segregated pathways within striatonigral projections, with the (dorsolateral) limb region of striatum projecting to ventromedial SNr, and the (ventrolateral) orofacial region of striatum projecting to dorsointermediate SNr (Foster et al., 2021; Lee et al., 2020).

We estimated the location of SNr neurons using 4-shank silicon probes spanning the putative limb and orofacial regions (Figure 9a; n=510 units/11 animals/21 sessions; of which n=241 units/8 animals/12 sessions were from animals performing self-timed movements, n=269 units/3 animals/9 sessions for spontaneous movements). As described above, we performed permutation tests to identify units with significantly positive or negative signs of peri-movement modulation (p<0.01).

**Figure 9.**
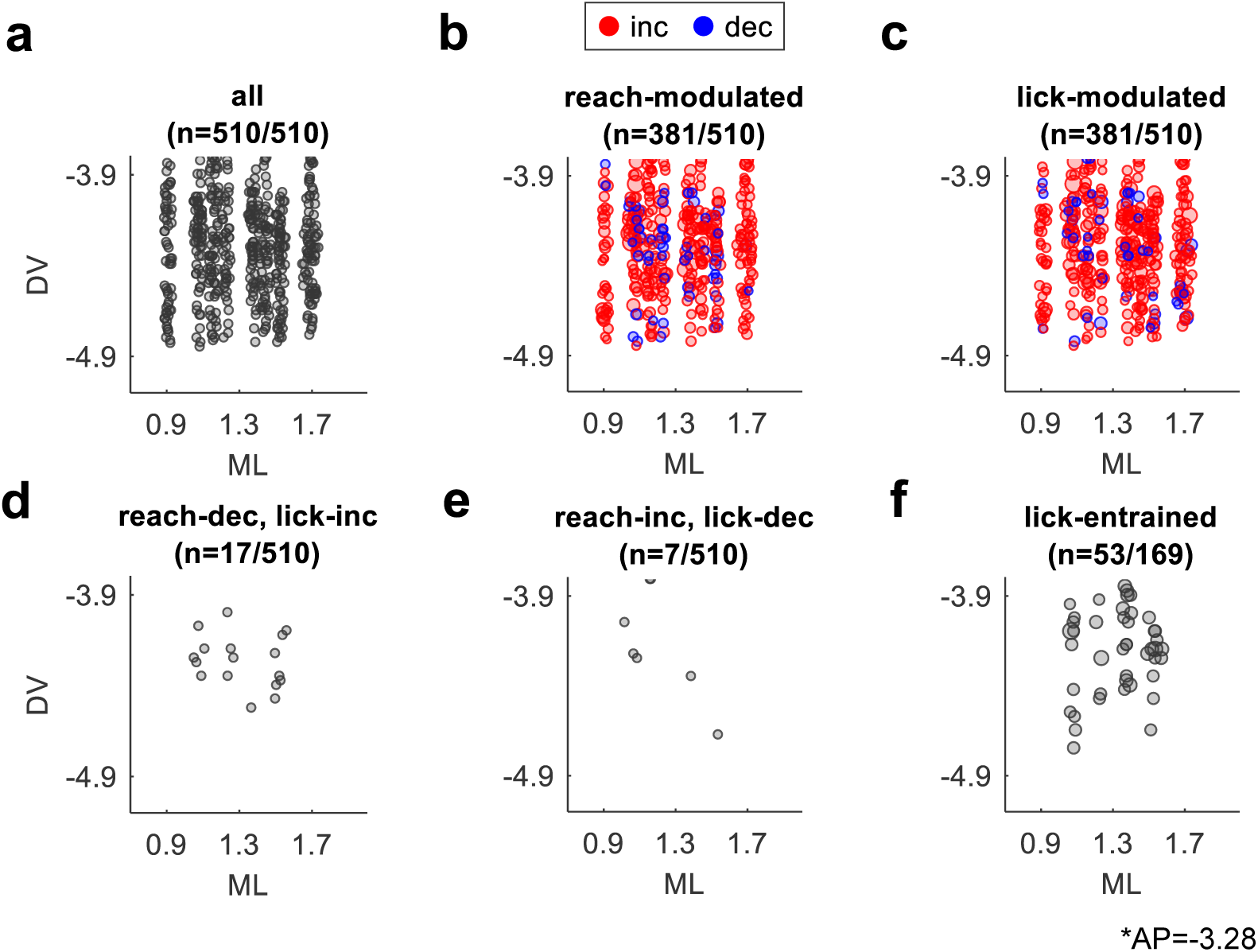
Estimated location of reach vs. lick modulated SNr units. **a.** Estimated location of recorded SNr units (n=510 units, 11 animals, 21 sessions). All coordinates are relative to Bregma. **b & c.** Location of SNr units significantly modulated during the reach (n=381) and lick (n=381) tasks, respectively. Red and blue circles indicate increase or decrease units, respectively. Radius of circles represent amplitude of movement-related modulation. **d.** SNr units which decreased spike rates during reach but increased during lick (n=17); or **e.** increased during reach but decreased during lick (n=7). **f.** Lick-entrained SNr units (n=45) from a subset of data (169 total SNr units, 8 animals, 12 sessions), these sessions were free of electrical artifacts in the peri-lick period. Coordinate estimates are given relative to Bregma (AP=-3.28, ML/DV as shown in figure).

Assuming that neurons that decrease their firing rate for a given movement type will disinhibit that movement, we would expect that *reach-decrease* and *lick-decrease* units would congregate in the ventromedial (limb) and dorsolateral (orofacial) portions of our recording sites, respectively. We visualized the approximate stereotaxic coordinates of 381 units significantly modulated during the reach task (Figure 9b) and 381 units modulated during the lick task (Figure 9c). Contrary to our prediction, we found no obvious pattern of segregation of reach- or lick-modulated units in the SNr. Furthermore, 24 SNr units with opposite signs of modulation for lick vs. reach (17 reach-decrease/lick increase and 7 reach-increase/lick-decrease; Figure 9d & e) were distributed across the four shanks. Lick-entrained neurons had been previously observed in the dorso-intermediate portion of SNr (Rossi et al., 2016), which preferentially receive input from ventrolateral (i.e. orofacial) striatum (Foster et al., 2021; Lee et al., 2020). However, 53 lick-entrained SNr units were evenly distributed throughout all four shanks of our recording probes (Figure 9f), including but not limited to the dorso-intermediate part of SNr. Thus, we did not find evidence for clear *functional* somatotopy within the SNr, in contrast to the segregated striatonigral circuits that have been defined by monosynaptic viral tracing.

## Discussion

We made three main findings. First, many SNr neurons modulated their firing rate before self-timed movements, with most neurons increasing rather than decreasing firing. Second, between different types of movement, the vast majority of SNr neurons had congruent signs of modulation, either increasing or decreasing; only a small minority of SNr neurons showed opposite signs of modulation. Third, for many neurons the modulation in firing rate began at least hundreds of milliseconds before the onset of self-timed and spontaneous movements. These observations suggest constraints – and raise new questions – about the role of the basal ganglia in movement initiation and action selection.

### Relating our findings to the action-selection model

Since the 1980s, researchers have observed a paradoxical pattern of activity in basal ganglia output neurons during behavior. First, many studies have reported that the majority of SNr and GPi neurons increase rather than decrease their firing rates with movement. We likewise found the preponderance of SNr neurons that increased their activity leading up to movement, in all the experimental contexts that we tested. SNr and GPi output neurons are GABAergic and tonically active; why would the basal ganglia relay even more inhibition to downstream targets around the start of movement? Second, most GPi neurons were previously reported to activate concurrently with or after the onset of EMG activity, seemingly too late to play a role in movement initiation. Yet hypo- and hyper-kinetic movement disorders suggest a role for the basal ganglia in movement initiation. How can the basal ganglia contribute to movement initiation if their output modulation does not precede movement?

To reconcile these observations, more than 30 years ago Mink and Thach proposed that basal ganglia output neurons could be involved in *selecting* rather than initiating movements (Mink & Thach, 1991a; Mink & Thach, 1993). They hypothesized that the majority of pallidal output neurons increase firing to inhibit unwanted actions, whereas a minority decrease firing to disinhibit the specific action desired at that moment. They further speculated that structured input from the antagonistic striatal direct and indirect pathways could shape antagonistic neuronal responses in the GPi/SNr (Mink, 1996). Indeed, subsequent findings in the striatum, such as co-activation of striatal dSPNs and iSPNs during movement, have been interpreted in this light (reviewed in Bariselli et al., 2019)

We set out to test this model by recording from SNr projection neurons in animals trained in self-timed or spontaneous movement tasks. We chose these behaviors based on clinical observations that Parkinson’s patients have more difficulty generating self-initiated movements compared to externally cued movements. We specifically tested an implicit prediction of the model, that SNr neurons should show a *reversal* in the *sign* of their modulation for different movements. That is, neurons that decrease firing to disinhibit a specific movement, *X,* should increase their firing to suppress movement *X* when the animals makes a different movement. However, among the hundreds of SNr neurons recorded in our various experiments, only a small fraction showed this sign-reversal phenomenon. This was the case whether the animals alternated between reach and lick movements (9% of neurons), or reached to medial vs. lateral targets (0% of neurons). Rather, the vast majority of SNr units evinced sign-congruent responses for the different types of movements, either increasing or decreasing firing for *both* movements. This was also true for SNr neurons that project to the SC, and area known to be involved in movement.

### Potential explanations for the paucity of sign-inverting SNr neurons

At first glance, our findings seem largely incompatible with a simple action-selection model in which SNr neurons select/suppress actions by inverting the sign of their peri-movement responses. However, there are several hypothetical scenarios whereby we could have missed sign-inverting neuronal responses or over-estimated the prevalence of sign-congruent neuronal responses. Consider first the neurons that decreased firing for both types of movements. It is possible that both tasks engaged a common movement that was not explicitly instructed yet still necessary for both tasks, such as a postural adjustment of the body. According to the action-selection model, if a neuron were selective for that common movement, it should decrease firing to disinhibit that movement in both tasks. Second, consider the SNr neurons that increased firing for both movements, e.g., reach and lick. These neurons might be selective for a third, unrelated movement that is not involved in either licking or forepaw reaching, such as movement of the hind-paw or tail. By increasing firing during licking *and* forepaw reaching, the congruent increase neurons would thus suppress that undesired movement during both tasks. Under these two scenarios, our finding of congruent (non-inverting) modulation could still be compatible with the action-selection hypothesis. In this view, had we tested a broader range of movements, we might have found more neurons with sign reversals.

However, there are several arguments against these hypothetical scenarios. First, licking and reaching are distinct mechanically, and would thus likely involve different compensatory postural adjustments. We also video-tracked other parts of the body (e.g., spine) during licking and reaching and did not observe any systematic common movements. Thus, we did not uncover evidence for a “third”, common movement that might otherwise explain the responses of congruent decrease neurons.

Second, these scenarios would imply that the SNr neurons with congruent responses – the vast majority in our study – were selective for some (unknown) movements *other than* reaching or licking. However, representations of the contralateral forearm/paw and orofacial are expansive in somatotopic motor maps in rodents (Hintiryan et al., 2016), and we deliberately targeted electrode penetrations to sample from areas of the SNr reported to represent forepaw and orofacial movements (Foster et al., 2021; Lee et al., 2020). Furthermore, we were able to identify SNr neurons projecting to the arm or orofacial regions of motor SC where artificial stimulation (or inhibition) was known to trigger (or suppress) arm or orofacial movements. These SC-projecting SNr neurons still overwhelmingly increased firing for licking and reaching, presumably sending additional inhibition to (rather than disinhibiting) motor SC. We also examined a broader range of movements that were associated with brief dips in SNr firing rates outside the context of behavioral tasks, but the incidence of neurons with transient dips that were specific for *one* type of movement did not exceed that expected from chance.

A recent study reported that most SNr neurons exhibited decreased firing during specific phases of reaching, retracting and chewing (Falasconi et al., 2025). We thus also examined SNr modulation during additional phases of movement such as arm extension vs. retraction, as well as the start vs. end of lick bouts. While a small percentage SNr neurons that decreased firing for one specific movement phase, a much larger population increased firing for all phases, and an additional fraction decreased firing for more than one or even all phases. Thus, movement-specific or behavioral-phase-specific decreases in firing were uncommon in our extensive data set, in contrast to the results of Falasconi et al. This difference is unlikely to be due to our sampling different populations of SNr neurons, because our recording sites covered areas that likely overlapped with the sampled regions in Falasconi et al. It is also possible that the transient movement-specific dips in Falasconi et al. were idiosyncratic to their precision reach-grasp-consume task – although specific dips in firing were also uncommon in our 2-direction reach task, for which the animals made more precise forelimb movements.

Importantly, in all the above analyses of peri-movement modulation, we normalized the spiking rates of each SNr neuron to its baseline spike rate, averaged over a period 2-4 s before the first self-timed reach or lick. Our animals typically “settled down” and remained stationary during this period; thus our baseline firing rate reflects SNr activity with little or no ongoing movement. If we had included spiking data from the peri-movement period when spike rates were often higher than baseline, our estimate of the baseline spike rate would have been inflated, and following baseline subtraction, we thus could have mistakenly overestimated the number of SNr neurons with opposite signs of modulation during different phases of movement. This issue could be exacerbated in freely-moving animals (as in Falasconi et al., 2025, where spurious movements before operant trials could contribute to a biased estimate of baseline firing rates. Lastly, we carefully excluded or filtered out electrical artifacts generated by the servo motor/touch detector/lick detector, because we found that these artifacts sometimes occluded spike detection during the peri-movement period.

Apropos, many neurons in our sample showed clear cycle-by-cycle entrainment in their pattern of firing during the repetitive lick bouts on rewarded trials. These neurons could be involved in protruding or retracting the tongue, or in modulating other periodic orofacial movements intrinsic to collecting and consuming the liquid (e.g., jaw movement, swallowing, etc.). By any definition, this pattern of neuronal activity is “lick-related” – yet the vast majority of these lick-related neurons still showed congruent activity between licking and reaching, either increasing or decreasing their firing rate before both types of movement. The reach task involved a sequence of reaching first and then licking after a delay, yet we found no evidence that the pre-movement ramping activity on reach trials was due to the upcoming lick; on the contrary, activity related to licking was delayed commensurate with the ∼1s delay between reaching and licking on those trials (Figure 6k). Under the action-selection hypothesis, we should not have observed lick-entrainment among the “congruent-increase” neurons, because they would have encoded some other movement(s) not involved in *either* reaching or licking; nonetheless, the vast majority of lick-entrained neurons showed congruent modulation between the lick and reach tasks (Figure 6j).

Given these contradictions, it is difficult to square our main findings with the simple action-selection model, whereby particular movements are selected by specific decreases in firing of SNr neurons. While it is still possible that the small minority of SNr neurons with sign-reversing responses might mediate antagonistic action selection in the SNr, the majority of SNr neurons could play a fundamentally different function during movement.

### Other potential roles for SNr output neurons

If most SNr neurons do not contribute to an antagonistic circuit for action selection in the SNr, what is their function? Thus far, we have emphasized that the *sign* of neural modulation was often congruent between movement types, but the *amplitude* of that modulation often differed. In fact, the majority of SNr neurons exhibited significantly different firing rates for forepaw reaching vs. licking (although surprisingly not for medial vs. lateral reaches made with the same forepaw). Presumably, downstream targets of the SNr could similarly decode movement type from SNr output. It is possible that these signals could then be harnessed for an antagonistic action-selection mechanism, but that antagonism would be played out downstream from the SNr, for example in the brainstem (Lee & Sabatini, 2021).

Notwithstanding, more generally, what other role(s) could be played by the majority of SNr neurons with congruent modulation between actions? It is possible that these neurons *always* decrease (or increase) their activity before *any* movement. In this view, the decrease neurons could nonspecifically disinhibit or gate downstream targets before any movement, with the specific movement dictated by brain structure(s) other than the SNr. Likewise, increase neurons could provide nonspecific inhibition for purposes other than action selection. For example, it has been suggested that cortical-basal ganglia-thalamic circuits act as large-scale feedback loops (Alexander et al., 1986); if so, increase neurons could play a role in stabilizing network output, similar to how negative feedback is commonly used in amplifier circuits to control and linearize output (Sanger, 2003). Feedback inhibition might also provide negative reafference to allow sensory systems to distinguish stimuli arising from external or self-derived sources (von Holst, 1954). There is also evidence in songbirds that pallidal spikes trigger precisely timed rebound excitation in postsynaptic thalamic neurons during singing behavior (Goldberg & Fee, 2012). In this view, increased pallidal output could precisely control the spiking pattern in target neurons, rather than just acting to inhibit firing. While these ideas are speculative, our findings suggest that, contrary to classic disinhibition or action-selection models, the SNr projects *more inhibition* around the time of movement, including to movement-related midbrain structures that are – at the same time – *increasing* their firing to drive movement. Future work will be needed to understand the role of this “paradoxical” movement-related inhibition (Goldberg et al., 2013).

### Timing of peri-movement modulation of SNr neurons

We found that pre-movement modulation in many SNr neurons preceded the actual self-timed movement by at least hundreds of ms, in the form of gradual ramping increases or decreases when averaged among trials for single units. The pre-movement activity preceded the earliest EMG activity and could not be attributed to an accelerating pattern of uninstructed jittery movements. We also found early pre-movement modulation in SNr neurons in a “spontaneous” movement task, with no explicit timing requirement. Previous work in monkeys had reported that GPi neurons were modulated late compared to the onset of movement, but those studies generally examined movements made as rapid reactions to external sensory cues; the late timing may have been an idiosyncrasy of rapid reactions. Using self-timed movements, we and others have observed modulation of basal ganglia activity well in advance of movement or EMG activity, in monkey putamen (Lee & Assad, 2003), mouse SNc (Hamilos et al., 2021), mouse striatum (Yang et al., 2024) and now mouse SNr. Thus, for self-timed movements, basal ganglia signals are present early enough to potentially influence movement initiation. Movements made in the absence of external triggers could require a gradual “revving” up of movement circuits that include the basal ganglia, perhaps as a component of a recurrent circuit with thalamus and cortex.

We did not examine the necessity of the SNr for self-timed or spontaneous movements. Lesions of pallidal output nuclei have little effect on reaction times for *cued* movements (Turner & Desmurget, 2010). However, our lab previously found that reversible optogenetic activation or inactivation of SNc dopamine neurons shifted the distribution of self-timed movements to earlier or later times, respectively (Hamilos et al., 2021). Given that the basal ganglia are the major recipient of dopaminergic output from the SNc, perturbations of the SNr may indeed affect self-timed or spontaneous movements. This will be important to examine in future experiments but may require methods to target specific functional cell types in the SNr.

### Functional vs. anatomical somatotopy

Somatotopic maps have been described in mouse cortex (Hintiryan et al., 2016) and striatum (Lee et al., 2020), and somatotopy has been inferred in the SNr by tracing monosynaptic striatonigral projections (Foster et al., 2021). However, we did not find evidence for *functional* somatotopy within the SNr in the context of lick vs. reach task, despite recording across the orofacial and forearm regions proposed by anatomical tracing. Rather, we observed a salt-and-pepper mixture of neurons that increased or decreased activity for licking or reaching, across a broad extent of SNr. Sign-reversing neurons, which may specify particular movements, were likewise broadly distributed. Even lick-entrained SNr neurons were not limited to the “lick region” suggested by anatomical tracing. The lack of clear functional somatotopy could arise from input to the SNr from the subthalamic nucleus (STN). STN input to the SNr/GPi is more spatially divergent than striatal input (Mink, 1996), and a recent study reported that STN neurons are more broadly tuned for multiple movement types (Wu et al., 2025). Regardless, the lack of clear functional organization might distract from more interesting organization at the level of genetically defined cell types. Single-cell RNA sequencing has revealed distinct cellular sub-populations within mouse SNr (Mendelsohn et al., 2024) and entopeduncular nucleus, the rodent homolog of the primate GPi (Wallace et al.), with different projection targets. An intriguing possibility is that the different neuronal responses that we observed in the SNr (i.e., sign-reversing, congruent increase and congruent decrease) could correspond to genetically distinct cell types. Understanding the inputs to and outputs from these neurons could help clarify their functional role. For example, do particular functionally defined neurons project differentially to brainstem or thalamus/cortex, and can they be distinguished based on patterns of input from the direct or indirect basal ganglia pathways? Addressing these questions will be essential to understanding the role of the basal ganglia output during normal behavior as well as in movement disorders.

## Methods

### Behavior tasks (self-timed reach/lick, directional reach, and spontaneous reach)

A total of 43 animals (22 female, 21 male; aged 2-12 months; C57BL/6J; Strain #000664; The Jackson Laboratory, Bar Harbor, Maine) were used in this study. All experiments and protocols were approved by the Harvard Institutional Animal Care and Use Committee and were conducted in accordance with the National Institutes of Health Guide for the Care and Use of Laboratory Animals. Animals were water-restricted 3 days prior to the first training session. Animals received ∼1mL of water or juice each day, adjusted on a daily basis to maintain 80% of their normal bodyweight.

For self-timed reaching (or self-timed licking) tasks, animals were head-fixed and placed on a raised metal platform. An aluminum touch bar was placed 0.5-1 cm in front of the animal, level with or slightly higher than the platform. The touch-bar and lick-spout were each connected to an electronic touch-detection circuit. Before start of a self-timed reach (or lick) trial, the touch-bar (or lick-spout) was deployed into range with a servo motor. When the touch-bar (or lick-spout) was fully deployed (800 ms traversal time), an audio cue (6,272 Hz tone; 100 ms duration) was played to indicate the start of the timing interval. If the first touch (or lick) was made ≥4 s after cue (correct trial), juice reward was dispensed via a solenoid valve connected to the lick spout; for reach trials, the lick spout was only deployed into licking range when a correct reach was made, effectively delaying reward delivery by 800 ms. After reward delivery, the touch bar and lick spout remains deployed until the end of trial (10 s after cue), at which time they were both retracted via servomotor. In rare cases where animals made a correct movement close to the end of trial, an extra 2 s were allowed for reward collection before retracting the bar and spout. If the first touch (or lick) was made <4 s after cue (incorrect trial), the touch bar (or lick spout) was immediately retracted, followed by a time-out period until end of trial (10 s after cue). If no movement was made during the 10 s trial period, the touch bar (or lick spout) was retracted at 10 s. In all cases (correct, incorrect, no-move) trials were 10 s in length, followed by a random intertrial interval (ITI) ranging between 3-10 s.

For the spontaneous reach task, a small touch-bar (0.4 cm wide, 0.8 cm tall) was placed in front of the contralateral paw. Touching the bar always resulted in its immediate retraction, followed by immediate redeployment (i.e. “cycling” the touch bar, 1400 ms total traversal time). The first reach in a session was always rewarded. Juice was delivered via an advancing lick spout. The lick spout was retracted at the end of the reward collection window (3000 ms, or up to 4000 ms if animals were still licking). The rewarded reach was followed by a random timeout period (drawn from exponential distribution, λ = 20 s; capped at 10 s to keep sessions reasonably short), during which reaches were not rewarded, and only resulted in “cycling” the touch bar. The first reach after the timeout period was rewarded. No external cue indicated the availability of reward. The exponential time-out period was designed to discourage timing behavior or rhythmic movements. Because many timeout periods were capped at 10 s, the distribution of inter-reach intervals for each behavior session was analyzed to ensure animals made spontaneous movements (exponential distribution) rather than timed movements every 10 seconds (Gaussian distribution).

The spontaneous lick task is similar to the spontaneous reach task, except for deploying/retracting the lick spout rather than the touch target. Some animals were trained to perform both spontaneous reach and lick tasks in the same session, where the task was switched every 15 rewarded trials.

For the bi-directional reach task, a small aluminum touch-bar (0.4 cm wide, 0.8 cm tall) was presented to the contralateral forepaw (determined relative to SNr recording site). Before the servomotor deployed the target into reaching range, a motorized linear translator first moved the target laterally to one of two pre-defined positions: 0.75 cm lateral to the resting position of the contralateral paw, or 0.5 cm medial to the contralateral paw. Target positions were chosen so that animals were forced to alter their movement trajectories. The target position changed every 15 rewarded trials. The bi-directional reach task is otherwise identical to the spontaneous reach task. (For the four-directional reach task, two additional targets were added: one in front of the contralateral paw and one in front of the ipsilateral paw. To reduce the complexity of training in the four-directional reach task, animals were rewarded for reaching immediately after the target was deployed. If animals did not move, the target was retracted at 10 seconds and redeployed after a random ITI.)

In all cases, the task flow was controlled in closed-loop with a Teensy microcontroller (PJRC) running a finite state machine program. Behaviorally relevant events (touch-bar contact, lick, cue, reward, etc.) were detected (if not generated) by the microcontroller and sent to the electrophysiology system as digital signals, as well as to a host PC as timestamped events. The host PC ran a custom MATLAB-based graphical user interface, which was used to send control signals (start/stop experiment, change task parameters, etc.), monitor task performance, and log behavioral data to disk.

### Surgery and histology

Mice were anaesthetized with isoflurane and head-fixed to a surgical stereotax using earbars. Fur was shaved from the skull, and the scalp disinfected with 70% ethanol and povidone-iodine. Zero pitch was achieved by leveling bregma and lambda points, and zero roll was achieved by leveling two points located 2mm left and right of midline. A lightweight titanium headpost was fixed to the skull with dental cement (C&B Metabond; Parkell, Edgewood, New York). The headpost was lowered and affixed to the skull while held with a custom fitting that ensured the headpost was horizontal in stereotaxic coordinate frame. Craniotomies for chronic or acute recordings were made with a stereotax-mounted drill and a 0.9 mm diameter drill bit. For chronic recording with microwire bundles, the moveable electrode wires were first covered with petroleum jelly, and the miniature microdrive was cemented to the skull and surrounded by a protective cone and a round cap (made of plastic printer film). For acute recordings, the craniotomy was made under anesthesia 2 hours before the first recording session and covered with a silicone sealant (Kwik-Cast; World Precision Instruments, Sarasota, Florida) between sessions.

Stereotactic coordinates for SNr recordings were 1) intermediate-lateral: −3.28 AP/±1.6 ML, used in chronic 32-channel microwire-bundle recording; 2) Intermediate-medial: −3.28 AP/±1.3 ML, for 4-shank probe recording, with the shanks spanning ±1.075 to ±1.525 ML (UCLA) or ±0.925 to ±1.675ML (Neuropixels). Depth estimates were made relative to dura, and then converted back to Bregma coordinates; putative SNr units were typically found between −3.8 DV and −5.0 DV. Units found above −3.8 DV were excluded.

Stereotaxic coordinates for injections in motor SC were: −3.5AP, −2.5DV, 1.2ML (intermediate) or 1.65ML (lateral). AAV5-Syn-CoChR-GFP (50 nL; 3.8×10^12^ vg/mL; UNC Vector Core, Chapel Hill, North Carolina) and AAV5-Syn-ChrimsonR-tdTomato (50 nL; 4.6×10^12^ vg/mL; UNC Vector Core) were injected in intermediate and lateral motor SC, respectively. In the reverse-injection control, the ML coordinates were reversed for CoChR and ChrimsonR, while the DV coordinates for intermediate/lateral SC were −2.2/-2.6, respectively, for better targeting.

Histology was performed by euthanizing the animal with pentobarbital sodium and perfusion with 4% PFA (w/v) in 0.1 M PB. Brain tissue was kept in 4% PFA for 24 hours and then in 30% sucrose for at least 24 hours before cutting. Tissue was embedded in Tissue-Tek OCT (Sakura Finetek USA, Torrance, California) and frozen at −24 °C and sectioned at 50-80 μm thickness. Histology was used to verify implant location of recording electrodes. For chronic recordings, wire bundle tracts were visible in tissue. For acute recordings, the probes were stained with DiI (Thermo Fisher, Waltham, Massachusetts) on the final recording session prior to histology.

### Extracellular recording and spike sorting

Initial single-unit recording in SNr was performed with gold-plated NiCr microwires (12 μm diameter; Sandvik, Palm Coast, Florida). In early recordings (2 animals) groups of 4 microwires were spun together in tetrode configuration; later recordings used a single bundle of 32 microwires, secured together by ultrathin-walled shrink-wrap (103-0402, 0.3 mm internal diameter; Nordson Medical, Westlake, Ohio). The wire tips extended ∼1 mm past the end of the tubing, and were allowed to splay out in the brain to sample a larger region. For ground, a single tungsten electrode was placed in the center of the bundle. Electrodes were connected to a 32-channel Omnetics connector via a custom electrode interface board (EIB) and gold pins (EIB pins; NeuraLynx, Bozeman, Montana). The EIB was mounted on a 3D-printed microdrive headstage, operated by a stainless-steel screw (0.500” length, M1.4-0.3 metric threads). Turning the screw one full turn corresponded to ∼300μm change in depth, allowing for granular control of electrode depth after implantation. During training, the tips of the wires were kept ∼3.8 mm below dura, roughly 300 μm above the substantia nigra. After the animals were fully trained, we progressively lowered the electrodes into SNr over multiple sessions. The microdrive was moved at least 24 hours before recording sessions to allow wires to fully settle into place. In addition to sampling different depths, moving the electrodes increased the yield compared to a fixed implant.

To better obtain positional information for recorded units, and to improve recording yield, we switched to acute stereotaxic recordings using 128-channel 4-shank silicon probes (model 128DN; Yang et al., 2020), and later 384-channel 4-shank Neuropixels 2.0 probes (Steinmetz et al., 2021). Probes were introduced through a 0.9 mm (1.5×1.5mm for Neuropixels) craniotomy (made on the first day of recording), with the mouse head-fixed using bilateral headpost holders that maintained the headpost (and thus the head) level with respect to the stereotaxic axes.

For acute recordings, two stainless steel ground wires were implanted in the brain and crimped within a single gold-plated connector pin. The craniotomy above the recording site was drilled open before the first recording session. The craniotomy was covered with silicone sealant (Kwik-Cast; World Precision Instruments, Saratosa, Florida) between subsequent recording sessions, and bathed in saline during recording. For all recordings, electrodes were gold plated to have 300-750 kΩ impedance.

Signals from extracellular recordings were collected using a Cerebus System (Blackrock Microsystems, Salt Lake City, Utah) or an RHD2000 System (Intan Technologies, Los Angeles, California); Neuropixels recordings were collected using SpikeGLX. Amplified electrode signals was sampled at 30,000 Hz and band-pass filtered (250Hz-7500Hz Butterworth) before storing to disk. Relevant behavior events were recorded as digital signals and stored as timestamps.

Spike-sorting was completed using custom software in MATLAB. The threshold for spike-detection was set as 2.5 SD of the signal (SD is estimated from MAD/0.6745): this usually captured spike waveforms as well as noise. Spike waveforms were separated from noise via k-means clustering, or by fitting a Gaussian mixture model, then visualized after dimensionality-reduction (from 31D to 3D) via principal component analysis. Because simple unsupervised clustering methods sometimes produced poor separation, we improved automated results by manual curation (removing electrical artefacts, splitting, merging, and adjusting cluster boundaries). We aimed to extract only high signal-to-noise single-unit spike trains from recorded data. Units with low signal-to-noise ratios were considered multi-unit activity and not included in analysis. Additionally, inter-spike intervals (ISI) smaller than 1.5 ms were at odds with the refractory period of spiking neurons. Therefore, units were classified as *single units* only if no more than 5% of interspike intervals were shorter than 1.5 ms. In cases where the same neuron was recorded on multiple adjacent channels, duplicate units were detected via cross-correlation analysis and removed. All presented neuronal data were from well-isolated single units.

In particular, we checked carefully that spike waveforms of “single units” remained uniform throughout self-timed trials, to ensure that any observed *changes* in firing rates were not due to spike-sorting artifacts, other units interfering, etc. In some cases electrode drift caused units to “disappear” halfway through recording sessions: these drifting units were excluded from analysis.

SNr neurons are tonically active, with typical spiking rates ranging from 20 to 100 spikes per second (sp/s). Dopamine neurons in the SNc and dorsal parts of SNr stereotypically exhibit 6-8 sp/s baseline firing and wider spike waveforms. Therefore, only single units with baseline firing rates >15 sp/s were considered putative SNr neurons.

When evaluating neural activity after the initial movement, we excluded channels with electrical artifacts generated by the servo motor used to move the touch-bar and lick-spout. Servo-motor related artifacts were rare and restricted to a few sessions. However, tongue contact with the lick spout often generated a large electrical artifact that resulted in a ringing signal that lasted up to 50 ms. We removed these ringing artifacts with an 800 Hz highpass filter before spike-detection and were able to recover lost spikes during the 10-50 ms period after spout-contact. We were not able to recover spikes in the [−10, 10] ms peri-spout-contact window, spike rates during these periods were either interpolated or omitted during trial-averaging (when applicable). We also excluded units whose spike waveform varied significantly as a function of time in the inter-lick period. Visual inspection of a random sample of raw vs. processed data revealed minimal distortion of the timing and waveform of spikes as a result of our processing pipeline.

### Identifying the sign and onset of peri-movement neural modulation

For pre-movement modulation in self-timed (reaching or licking) tasks, only trials in which the animal waited ≥2 s after cue were included, since shorter trials may include reactionary movements to the cue. A peri-event time histogram (PETH) was generated for each SNr unit by counting spikes in 100-ms (or 25-ms, for visualization) bins aligned to movement time (bar or spout contact for reach or lick tasks, respectively). Spike counts are divided by bin width and then averaged across trials to estimate spike rates. For comparing across multiple units, the PETH for each unit is normalized by subtracting the mean and dividing by the standard deviation estimated during the baseline window ([−4000 ms, −2000 ms] prior to movement). For calculating response onset latency, a smoothed PETH was generated by first counting spikes using 1-ms bins, then convolving with a Gaussian smoothing kernel (σ=100 ms, width=1000 ms), then averaging across trials.

In order to test whether putative SNr neurons were significantly modulated by movement, on a unit-by-unit basis, we first calculated its PETH (100-ms bins) aligned to the first movement (bar/spout contact) in each self-timed trial. We then compiled the distribution of its spiking rate in the *peri-movement window* ([−300, 0] ms before touchbar contact), as well as the distribution of spiking rates in the *baseline window* ([−4000, −2000] ms before touchbar contact). This yielded a distribution of peri-movement spiking rates (sample size = 3*K*, where *K* is the number of trials, and 3 is the number of 100-ms bins in the peri-movement window), as well as a distribution of baseline spiking rates (sample size = 20*K*, across 20 100-m bins). We then measured the difference between the means of the peri-movement vs. baseline distributions, using a permutation test to assess whether this difference is statistically significant, under the null hypothesis that spike rates in each of the 20*K* baseline bins and 3*K* pre-movement bins are independent and identically distributed (i.i.d.). Spike rates in baseline and pre-movement bins were shuffled 100,000 times. For each permutation, we re-sampled 20*K* ‘baseline’ bins and 3*K* ‘response’ bins and calculated the difference between their means. The 99% confidence interval for this difference-of-means was calculated from 100,000 permutations. A unit was classified as significantly movement-responsive if the observed difference-of-means fell outside the 99% confidence interval (i.e., permutation test; p<0.01).

A similar permutation test was performed for 3 subsequent phases of movement in rewarded trails (“first lick”, “last lick”, and “bar release”). We chose a [−300, 300] ms peri-movement window (relative to the first spout contact, last spout separation, and bar release, respectively), compared to a baseline window [−4000, −2000] ms relative to bar-contact in reach trials. Permuting 20*K*_reach_ baseline bins and 6*K* peri-movement bins (100,000 permutations, where *K*_reach_ is number of self-timed reach trials used for baseline, *K* is the number of repeats for the movement-in-question).

For responsive SNr units, we then analyzed their response latency (i.e., onset time) using the kernel-smoothed PETH previously described. Looking back from movement time, response onset was defined as when the normalized, smoothed spike rates (sampled at 1 ms resolution) first exhibited 25 ms of baseline activity (<0.25 SD) followed by 50 ms of above-baseline (or below-baseline) activity (≥0.25 SD).

### Video tracking and movement onset detection

Bilateral video recordings of animal movement were collected using webcams at ∼30 frames per second. IR LEDs are used to illuminate the chamber during recording. Video streaming and logging was controlled via MATLAB (Image Acquisition Toolbox). A timestamp is registered by MATLAB on every 10th frame, which is used to align video and spiking data. DeepLapCut (Mathis et al., 2018) was used to train a neural network (ResNet50) to identify the location of relevant body parts (front and rear paws, nose, tail, and the midpoint of the spine). Model-predicted positions with low likelihoods (<95%) were discarded, because these were typically erroneous predictions due to occlusion of body parts.

To compare movement onset times against neural onset times, video frames were aligned with electrophysiology data (made possible by saving the timestamp of every 10th video frame, as well as the timestamp of the first electrophysiology sample). Accurate timing alignment was verified by having a LED visible to the camera turn on at the start of each trial, while the LED gating signal was recorded as a digital input on the spike-acquisition signal.

Electrically detected touchbar contact times were used to determine the end of arm reaches. Movement onset (forepaw lift) time was estimated from video-derived trajectories of the contralateral forepaw (resampled at 25-ms intervals), by tracing backwards from end-of-reach until the contralateral forepaw was observed at its resting position for 25 ms followed by 50 ms away from resting position (≥0.25 SD). Automatically detected movement-onset times as well as bar-contact times have been validated by manual inspection of sample trials.

In one animal (3 sessions), a pair of EMG wires were implanted in the neck muscles. The EMG signal were amplified (x1000 gain) and filtered (50-500 Hz band-pass) using an amplifier (Model 1700; A-M Systems, Sequim, Washington), half-wave rectified and recorded at 30,000 Hz (Intan RHD2000 evaluation board), then smoothed (33.3ms moving-average) and normalized (robust z-score) in MATLAB. To detect EMG onset, the smoothed, normalized EMG signal was resampled at 25-ms intervals, and EMG onset was determined by tracing backwards to the first time EMG exceeded 0.25 SD for two consecutive 25-ms samples.

### Generalized linear model for decoding SNr unit activity

For each SNr units, a series of nested GLMs were fitted to predict its spike rate time course. 4 categories of predictors were iteratively added to the model, with the goal of estimating whether model performance had been improved by including more predictors:

1. The *cue* predictor was generated by converting discrete trial-start events into a continuous time series across the session, using a set of 13 overlapping cosine kernels (width 400 ms, temporal offsets ranging from 0 to ±600 ms). Negative latency offsets were included because the servo motors engage 800 ms before trial start, a potentially salient cue for behaving animals.
2. The *velocity* predictors were generated from video analysis of body parts. Position traces detected by neuron networks were converted into 2D velocity traces, then normalized by the mean and SD (i.e., z-scoring). For bilateral videos, the x and y axes respectively point to anterior and dorsal side of the animal. Body parts visible to both cameras (nose, spine, tail) were averaged between cameras after normalization. Normalized velocity signals were then convolved with a series of 9 cosine kernels (width 100 ms, temporal offsets 0 to ±200 ms). Latency offsets of up to 200ms covered the range of neural response times previously observed in monkey SNr and GPi.
3. The *ramp* predictors were a series of linear ramps (0 to 1) terminating at touchbar-contact times; 19 such ramps starting 100-2000 ms prior to touchbar contact were added to the model. They were timing-invariant in the sense that these ramps did not begin at cue (when the animal begins timing) but were rather aligned to the movement time. We added the ramp predictors *after* the velocity predictors in the nested GLM approach: if the velocity predictors already explained the ramping activity, then adding the ramp predictors should not significantly improve model performance.
4. The *trial-progression* predictors were signals that increased linearly from 0 to 1 from trial-start to touchbar contact. These predictors capture elapsed time from cue up to the moment of movement.

GLMs were fit using iteratively reweighted least squares to find the maximum likelihood estimate for model parameters (Statistics and Machine Learning Toolbox, MATLAB). Five-fold cross validation was employed to reduce overfitting. Model performance was evaluated by variance explained (R^2^) and Akaike Information Criterion (AIC), calculating the additional variance explained (ΔR^2^) as well as the reduction of Akaike Information Criterion (ΔAIC) after adding more predictors to the model. Fitted models were used to predict single-trial spiking rate traces, which were then averaged across trials and compared to true observations.

### Linear discriminant analysis for decoding SNr population activity

Population SNr activity was collected during the peri-movement period for reach vs. lick trials ([−0.3, 0] s relative to bar/spout contact) and during the two-target lateral vs. medial reach trials ([−0.1, 0.2] s relative to reach onset). To ensure population size, only sessions with ≥10 recorded SNr units were included (10 animals/13 session for reach vs. lick; 3 animals/7 sessions for lateral vs. medial reach). Peri-movement spiking activity for each SNr unit was normalized to the mean and SD of its baseline spike rates (estimated during baseline window: [−4, −2] s before movement). For each session, normalized peri-movement activity as well as baseline activity were used to train a decoding model (linear discriminant analysis, or LDA) by fitting a Gaussian mixture model (assuming equal variance for all 3 states: lick/reach/baseline; or lateral-reach/medial-reach/baseline). We assumed empirical prior probabilities proportional to the number of observed trials of each type (490 reach vs. 680 lick trials vs. 1170 baseline periods; 351 lateral reach trials vs. 424 medial reach trials vs. 775 baseline periods). We implemented five-fold cross-validation to reduce overfitting. Models trained on the peri-movement window ([−0.3, 0] s or [−0.1, 0.2] s) were used to decode population neural activity from [−4, 2] s relative to movement; yielding model-predicted probabilities for each state: p(reach), p(lick) and p(baseline); or p(lateral), p(medial) and p(baseline) for the reach vs. lick task and two-target reach task, respectively. For each time point *t* (relative to movement time), we then took the difference in predicted probabilities: Δ*p*(*t*) = *p*(reach|*t*) - *p*(lick|*t*), or Δ*p*(*t*) = *p*(lateral|*t*) - *p*(medial|*t*) as a one-dimensional representation of the time-course of SNr population activity. Model predictions are averaged across true reach trials vs. true lick trials (or true lateral-reach trials vs. true medial-reach trials). For visualization, we then averaged decoded population activity across sessions (weighted by trial count). This grand average is representative of each individual session upon inspection.

To examine the significance of model-decoded activity, we also trained null models using shuffled training data where the lick/reach (or lateral/medial reach) trial labels were randomly permuted. 10,000 permutations were performed, each time generating a null model used to decode population activity. As expected, the null models were not able to decode SNr activity. The 99% confidence interval (i.e., 0.5% and 99.5% percentiles) for Δ*p*(*t*) was calculated amongst the 10,000 null models.

### Detecting differences across PETHs grouped by movement time

Self-timed reach trials were divided into 4 groups according to movement time (2-4s, 4-6s, 6-8s, and 8-10s; i.e., how long animals waited after the start cue before touching the bar; trials shorter than 2s were considered acute reactions to the cue and excluded from analysis). Thus, for each SNr unit, PETHs were calculated for 4 groups of trials: where *x*_*i*_(*t*) is spike rate averaged across all trials in group *i*, expressed as a function of time relative to bar contact *t* (where −2 s ≤ *t* < 0, discretized at 0.1 s resolution). 6 pairwise Euclidean distances were calculated among 4 PETHs: *d*_*i*,*j*_ = ∑_*t*_(*x*_*i*_(*t*) - *x*_*j*_(*t*)). Distances were only calculated in the window [−2, 0] s before bar contact where the majority of pre-movement activity was observed. We took 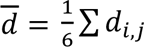, the mean across the 6 Euclidean distances, as a measurement of the dissimilarity between ramping neural activity across different movement times. Under the null hypothesis that there is no difference between groups (*d* = 0), we randomly shuffled trial identities before grouping and calculating the 4 PETHs, and generated a 99% confidence interval for *d*. The minority (30/641) of SNr units whose *d* fell outside the confidence interval were potentially timing-dependent and selected for further inspection.

### Dip/rise-triggered average kinematics

First, transient dips (or rises) in firing rate were detected for each putative SNr neuron throughout the session (n=1,225 units, 39 sessions, 6 animals). Spike rate was estimated from spike counts binned at 1-ms resolution before smoothing with a Gaussian kernel (SD=75 ms, width=500 ms), and z-scored relative to its whole-session mean and SD. The threshold for dips (or rises) were chosen as the 25^th^ (75^th^) percentile of all negative (positive) z-scored spike rates. A dip was detected when spike rate continuously exceeded threshold for 200-600 ms but did not exceed threshold for 100 ms before the onset, nor after the offset. Movement kinematics were aligned to the onset and averaged across dips or rises (i.e., dip-triggered averages or rise-triggered averages).

For movement kinematics, cameras were installed on the left and right side of the animal, recoding at 30 fps. DeepLabCut was used to track the position of the jaw, left hand, right hand, and the midpoint of the spine. Position was smoothed with a Gaussian kernel (SD=133 ms, width=667 ms) and then z-scored relative to its whole-session mean and SD. The left hand and the right hand were tracked with the left and right camera, respectively. The tongue, jaw and spine were tracked with both cameras and averaged across cameras. We focused on the DV position of the jaw and spine, and the AP position of the hands. Tongue protrusion was detected when DeepLabCut reported ≥50% likelihood of tongue tracking on either the left or right camera. Tongue-protrusion probability is then smoothed with a Gaussian kernel (SD=33 ms, width=167 ms).

All 836,887 dips and 777,994 rises (detected from 1,225 SNr units, 6 animals, 39 sessions, recorded acutely with Neuropixels 2.0 probes) were clustered (k-means) based on 5 features (jaw, tongue, left hand, right hand, and spine) into 7 classes (1 - “no movement on average”, 2 - “jaw open”, 3 - “jaw close”, 4 - “left hand forward”, 5 - “left hand back”, 6 - “right hand forward”, and 7 - “right hand back”). For each SNr unit, dip/rise-triggered kinematics for the jaw (tongue was redundant/omitted), left hand, right hand, and spine were averaged within each of the 7 clusters. For each cluster (n=7), its dip/rise-triggered average movement profile for each body part (n=4) was deemed significant if the observed change in position [−300, 0] ms before dip-onset vs. [0, 300] ms after dip-onset was higher than chance (p<0.05 with Bonferroni correction for 7×4=28 comparisons; i.e. p<0.0018). Chance level for dip/rise-triggered averages was estimated for each SNr unit by bootstrapping a null distribution (10,000 repetitions), each time randomly sampling *n*_*k*_ 600-ms intervals throughout the session (*n*_*k*_: number of dips/rises in cluster *k* for that SNr unit), then calculating the change in position across two halves of the 600-ms interval.

For each SNr unit, we excluded clusters of dips (rises) with <10 instances and clusters contaminated by unrelated movements (i.e., significant hand movements in the jaw-movement clusters; or vice versa). We tallied the number of SNr units containing the following 5 types of significant uncontaminated dip(rise)-triggered movements: jaw (either open or close), left hand forward, left hand back, right hand forward or right hand back. To estimate the chance level (i.e., null distribution) for this analysis, we performed a second bootstrap analysis (1,000 repetitions): each time 1) randomly sampling *n*_*k*_ 600-ms intervals, 2) calculated the average change in kinematics, then 3) estimating significance of said kinematics (mean ± 3SD for hands, mean ± 1.5SD for jaw and spine were considered significant movements; thresholds were chosen to minimize error rates matching the results of first bootstrap mentioned above; since performing 1,000 × 10,000 bootstraps would have been computationally expensive).

### Detection of lick-entrained units using circular statistics

We used circular statistics to determine if the spiking rates of SNr units fluctuated in sync with bouts of rhythmic licking. The interval between two adjacent licks (spout-contact) were evenly divided into 30 bins to generate a circular PETH, where each bin in the PETH represents the average spike rate during a certain phase (0-2π) of the lick. Interlick intervals longer than 200 ms (non-rhythmic licking) or shorter than 50 ms (likely due to lick-detector artifact) were not considered. For each unit, its circular PETH was expressed as 30 vectors in polar coordinates: (phase, spike rate). Averaging across these 30 vectors in Euclidean space yielded a “phase-selectivity vector”, whose magnitude should be zero for units with perfectly stationary spike rates. A non-zero magnitude indicated increased or decreased spike rates during certain phases of licking (i.e., lick-entrained). Therefore, we tested whether the average vector had a significantly non-zero magnitude using a permutation test as follows: 1) shuffle the phase of all 30 PETH vectors 10,000 times, each time calculating the magnitude of the average vector; 2) check whether the “observed” average vector magnitude falls within the top 1% of the distribution of “shuffled” vector magnitudes (i.e., p < 0.01).

While lick-entrainment was detected using lick-pairs, we tried to verify whether entrainment was present in longer lick bouts. To that end we visualized lick-entrainment by plotting an extended PETH spanning 5 consecutive licks. Longer lick trains were split into five-lick bouts (whereas shorter lick bouts were discarded); all five-lick bouts in a session were aligned by phase (0-8π, with multiples of 2π denoting spout contact) to generate a PETH (as seen in Figure 6h) showing lick-entrainment.

Electrical artifacts were removed as best we can (see Extracellular recording and spike sorting), while spike rates [−10, 10] ms around the time of spout contact were interpolated from neighboring bins, since it was difficult to completely remove artifacts during this period. An additional control experiment (2 animals, 10 sessions) was conducted with accelerometer/camera-based lick detectors which did not generate electrical artifacts. In the control experiment 25/249 SNr neurons were identified as lick-entrained, similar to the results shown in Figure 6g-k.

### Optogenetic stimulation of SC and opto-tagging of SC-projecting SNr neurons

To optogenetically activate SC neurons in head-fixed mice, we acutely implanted an optrode (Neuropixel 2.0 probe combined with a tapered optic fiber, 0.66NA, 200μm diameter, 1mm active length; OptogeniX, Arnesano, Italy). The optic fiber was lowered ∼2.7-3.0 mm below dura, at approximately −3.2AP, 1.4ML. Beams from a 470 nm laser and a 635 nm laser were combined and fed into the same patch cable (0.50NA, OptogeniX) before entering the optrode. 4 light input angles corresponding to 0.5NA, 0.33NA, 0.16NA, 0NA were used to vary the emission depth along the 1mm taper of the fiber (but combined by averaging during analysis). Laser power varied between 0.1, 0.5, 2, and 16 mW (measured at the tip of patch cable). Every combination of laser wavelength/power/duration were tested at least once (as a train of 10 pulses), in random order. Trains of 10 identical pulses (20ms or 100ms pulses at 1 Hz, or for 250 ms pulses at 0.5 Hz) were delivered at a time. Because opsin expression levels, fiber optic transmittance, and the precise placement of acutely-implanted optic fibers could be variable across animals, we tested combinations of laser powers and pulse duration, and chose the lowest light level/shortest pulse duration capable of reliably triggering movements. Movement generated by optogenetic stimulation of SC was monitored by video tracking of the jaw and the contralateral forepaw (DeepLabCut).

To opto-tag SNr neurons putatively projecting to SC, we acutely implanted the optrode in SNr (−3.28AP, 1.3ML, −4.8 DV). In between spontaneous lick/reach trials and at the end of the recording session, both the bar and spout were retracted, and a train of 10 identical 10-ms (or 20-ms) laser pulses were delivered at 10 Hz. Each train contained a different combination of laser wavelength/power (same as above). To obtain peri-stimulation spike rates at high temporal resolution, spike rates were estimated from the inverse of the inter-spike interval, as follows: 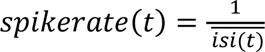; where for the *n*^th^ spike that occurred at *t*_*n*_, we define *isi*(*t*_*n*_) = *t*_*n*_ - *t*_*n*−1_ (i.e., time since previous spike); then using linear interpolation at 1-ms resolution between spikes, we defined 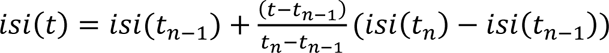 for *tn*−1 < *t* < *t*_n_; the *isi*(*t*) time series were aligned to laser onset and then averaged across trials to yield 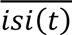. To calculate the “normalized stimulation response”, trial-averaged spike rates after each laser pulse were normalized to the mean and standard deviation of baseline spike rate ([−100, 0] ms prior to the onset of each laser pulse), units with significant increases in spike rate (>2SD) in the [5, 50] ms window after laser onset were considered opto-tagged. To determine the onset latency of stimulation response, we found the first time point after laser onset where normalized spike rate exceeded 2SD for two consecutive 1-ms samples. SNr units responding to >2mW 635nm light were considered ChrimsonR^+^, while SNr units responding to 0.5mW 470nm light but not >2mW 635nm light were considered CoChR^+^.

## Acknowledgements

We thank ML Andermann, B Sabatini, SR Datta, R Born, N Uchida, and M Wallace for research guidance; M Bukwich and C Starkweather for sharing electrode-making and implant techniques; E Sayed and T Wheatcroft for information on SC somatotopy; AE Hamilos, V Berezovskii, T LaFratta, J LeBlanc, O Mazor and P Gorelik for technical assistance.

---

**Figure 1—Figure Supplement 1.**
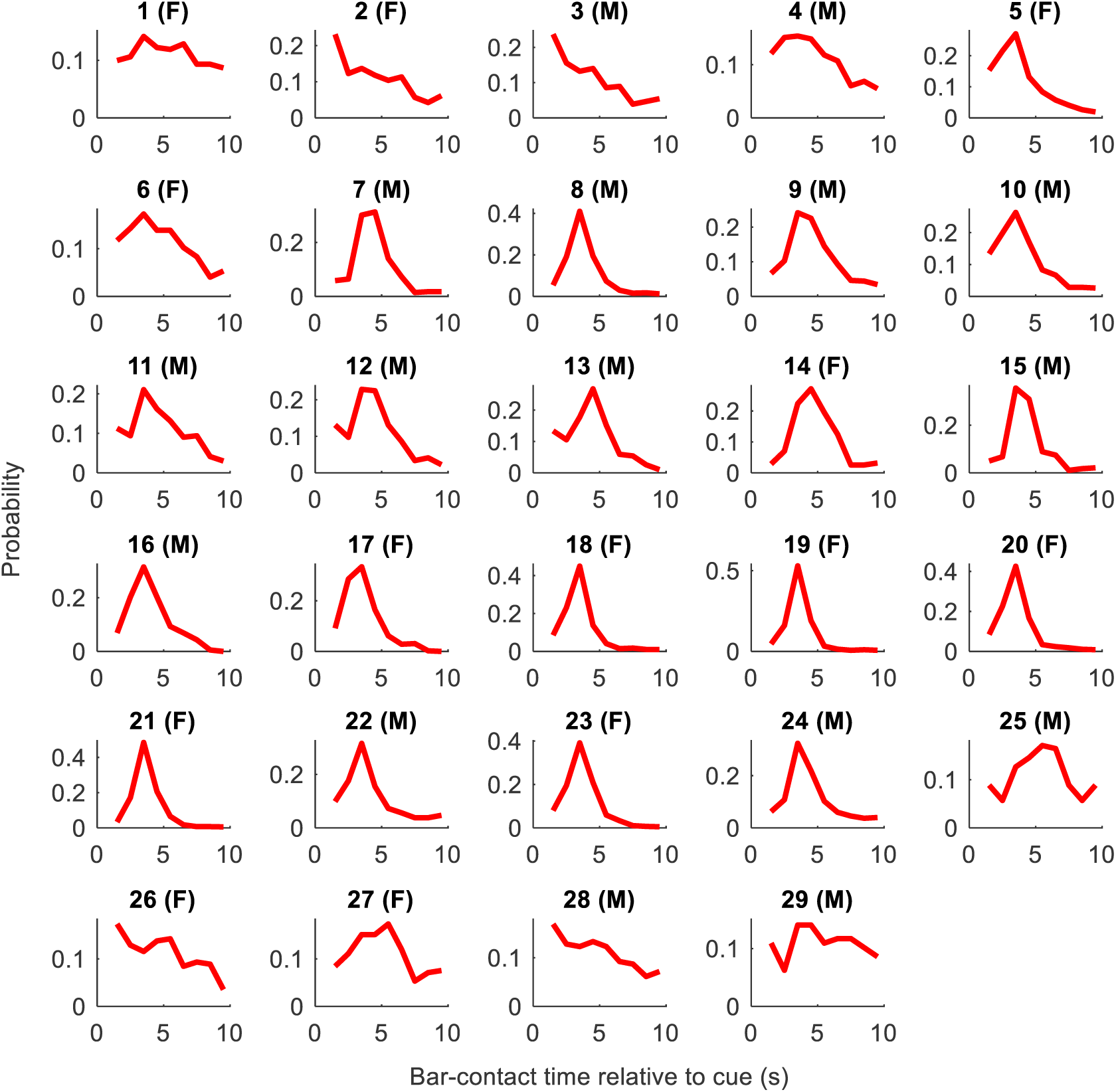
Movement time distribution for individual animals in the self-timed reach task. Movement-time histograms are generated for each of the 29 animals performing the self-timed reach task, aggregated across three best sessions after animals were fully trained. Reactive movements made before 1 s are not shown.

**Figure 2—Figure Supplement 1.**
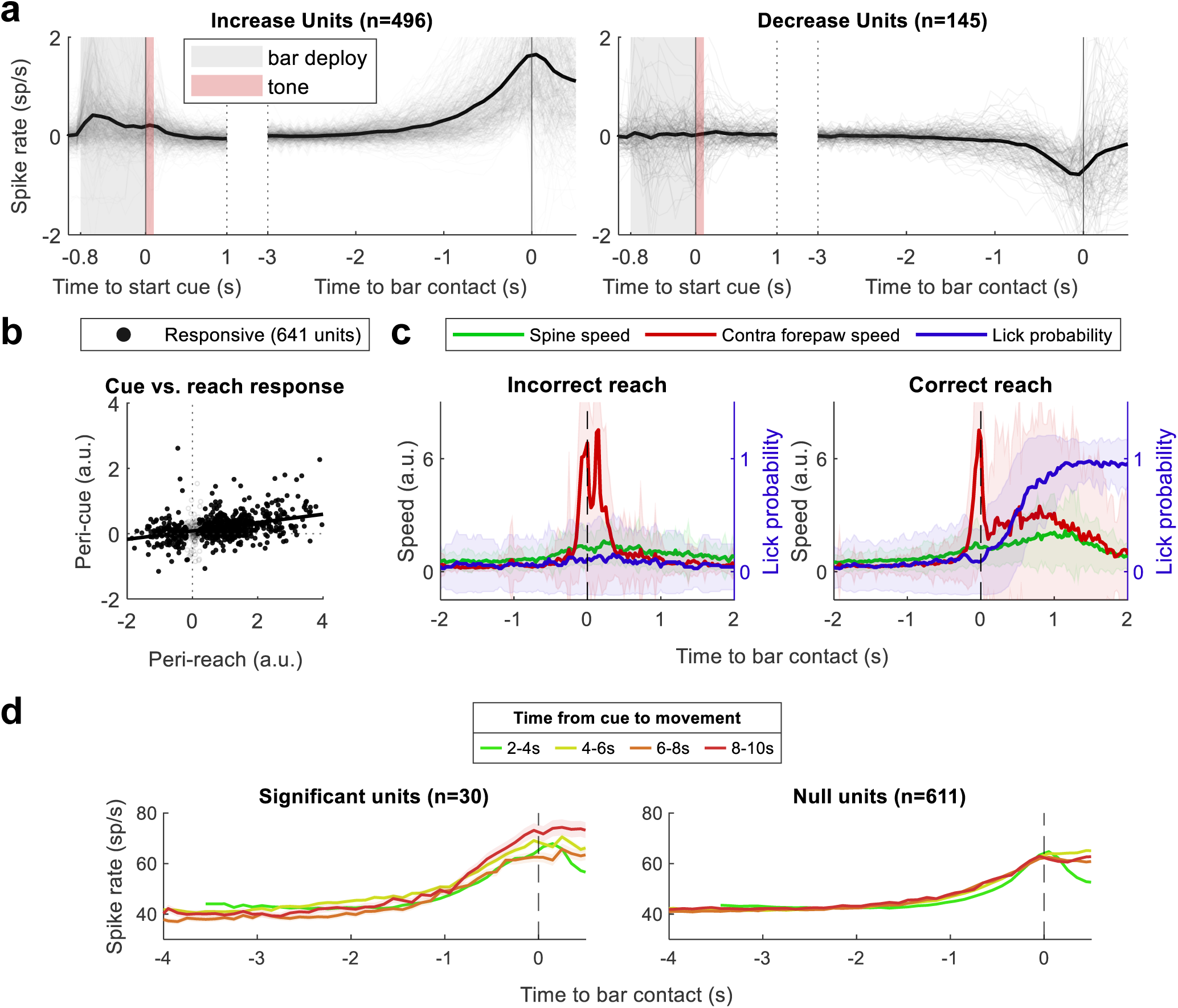
**a.** Cue-aligned and movement-aligned PETH normalized to baseline (left: 496 increase units; right: 145 decrease units). Dark black trace represents population average. Grey and red shaded areas represent bar deployment (800 ms) and tone (100 ms), respectively. **b.** Normalized peri-cue activity ([−0.8, 0.1] s relative to tone onset) vs. peri-reach activity ([−0.3, 0] s relative to bar contact). Black dots represent movement-modulated units (n=641), grey circles represent non-modulated units (n=143). Black line represents linear regression (R^2^=0.17). **c.** Trial-averaged normalized movement speed of the spine (green) and the contralateral forepaw (red), as well as tongue protrusion probability (blue), derived from video analysis. Plots are shown separately for incorrect/unrewarded (left, n=71) vs. correct/rewarded (right, n=185) reach trials. Shaded area represent standard deviation. **d.** Population average PETH during the self-timed reach task, grouped by movement time relative to the trial-start cue (as indicated by color). **Left**: population average across 30 SNr units whose pre-movement activity ([−2, 0] s before bar contact) differ significantly based on movement timing (p<0.01; permutation test); **Right**: population average across the remaining 611 SNr units whose pre-movement activity are similar regardless of movement timing.

**Figure 4—Figure Supplement 1.**
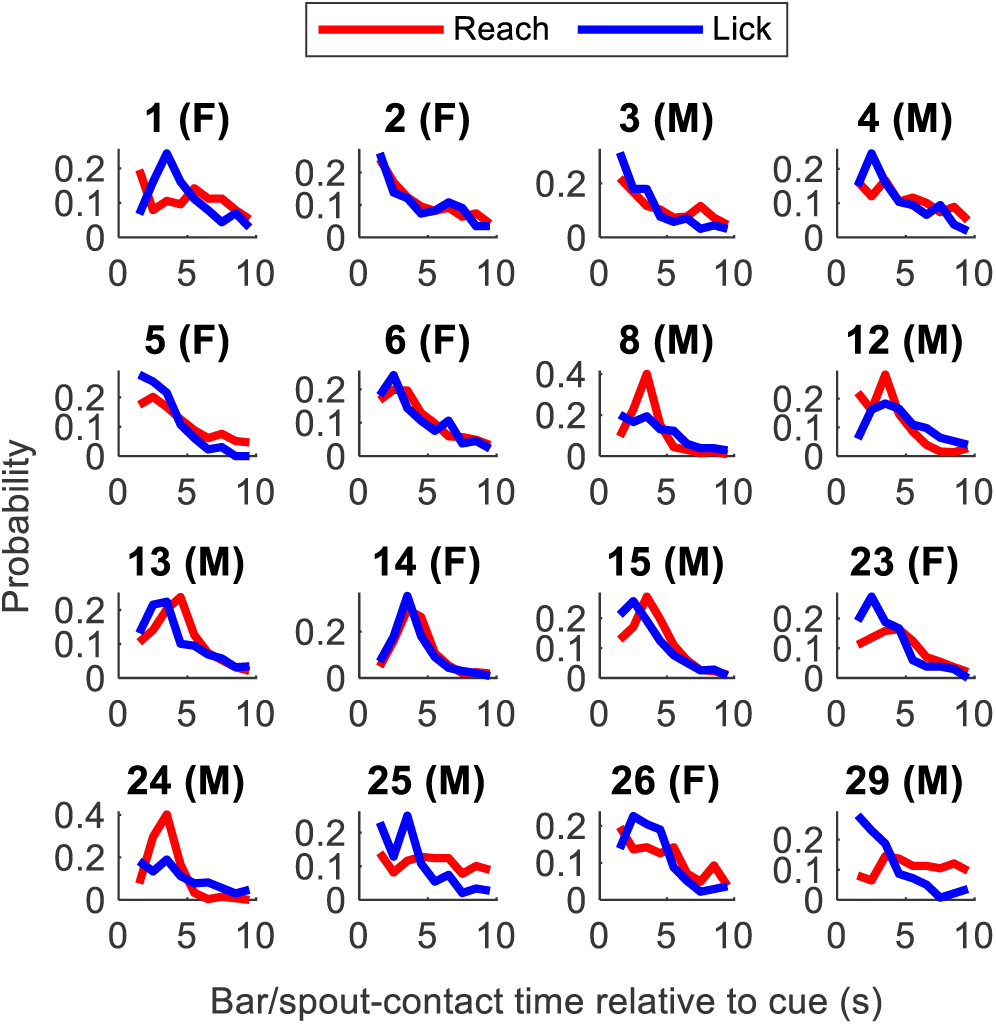
Movement time distribution for individual animals in the self-timed reach vs. lick task, for each of the 16 animals performing the self-timed reach vs. lick task, aggregated across three best sessions after animals were trained. Reactive movements made before 1 s are not shown. Reach times are shown in red; lick times are shown in blue.

**Figure 5—Figure Supplement 1.**
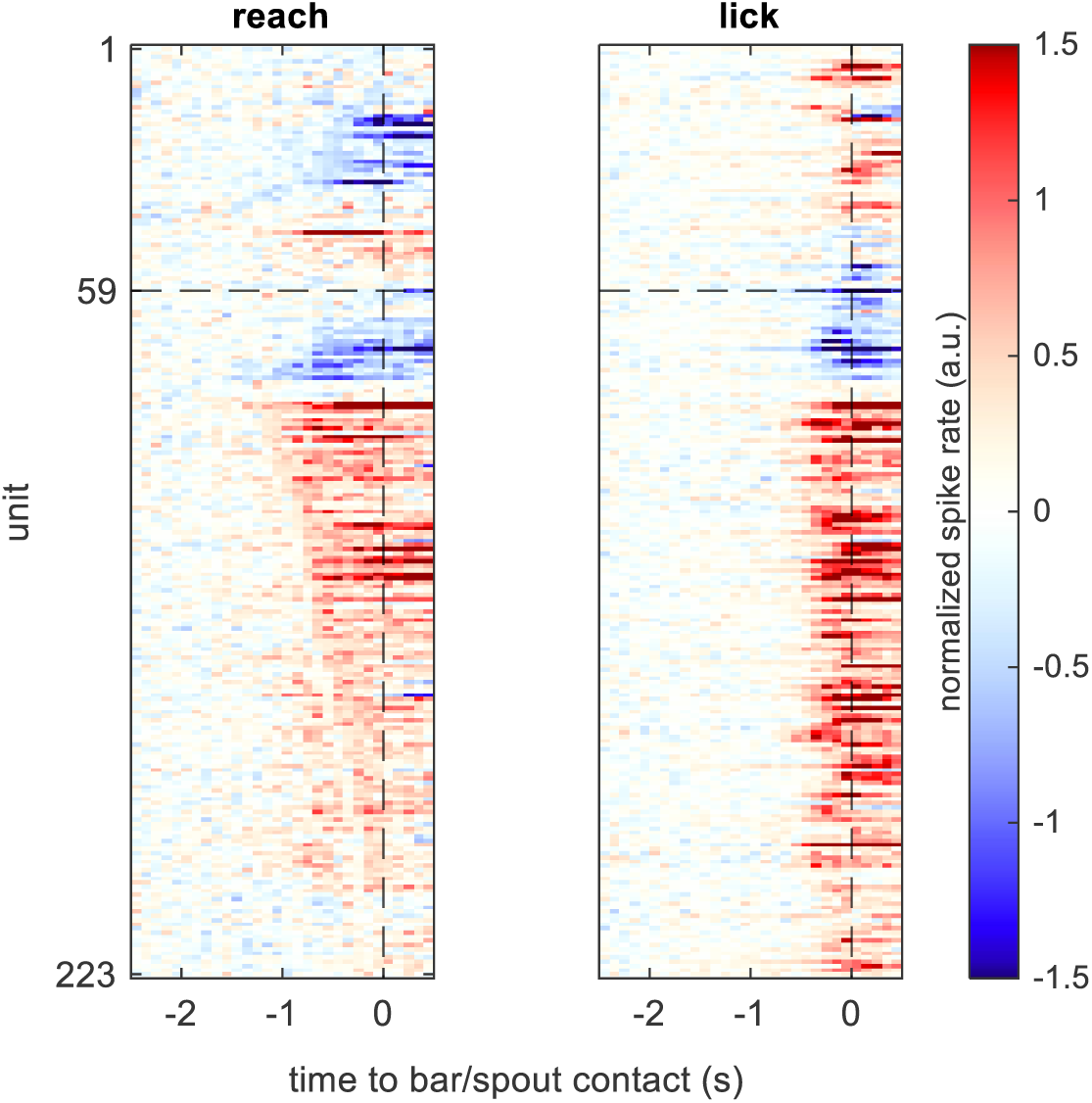
SNr modulation in the spontaneous reach vs. lick task. Heat map visualization of peri-reach (left) and peri-lick (right) PETHs for 223 SNr units arranged in the same order. The horizontal dashed line separates oppositely-modulated units (above) from congruent units (below). PETHs are normalized to baseline spike rates [−3, −1.5] s before movement. Only a subset of 223/1,225 SNr units with a minimum of 15 valid reach/lick trials (each) are visualized. Valid trials have inter-reach/lick intervals greater than 1 s.

**Figure 7—Figure Supplement 1.**
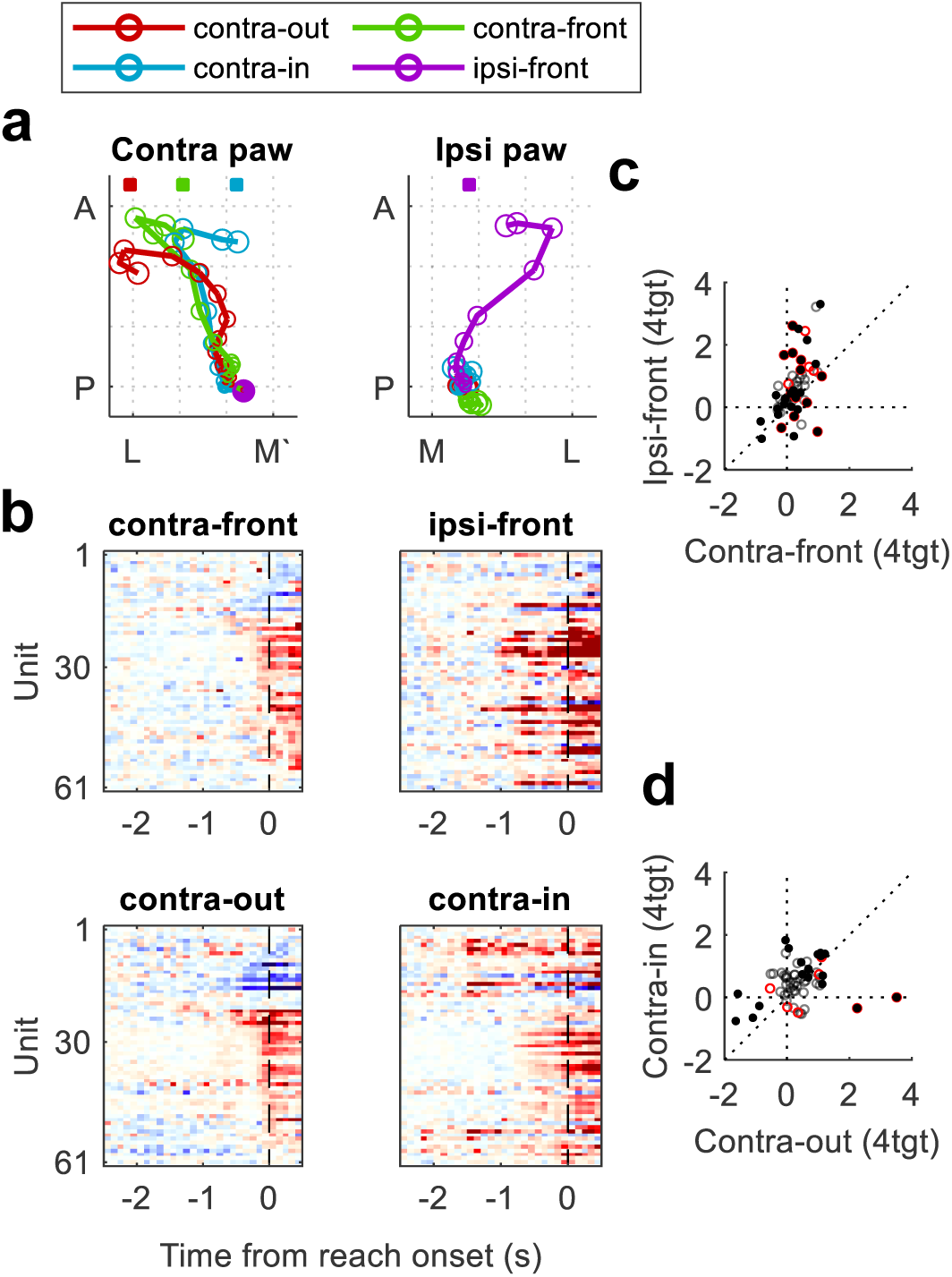
The four-direction forepaw reach task. We further trained two animals to perform the reach task with four possible target locations: lateral to, in front of, or medial to the contralateral paw, or in front of the ipsilateral paw (see Methods). We named these target locations “contra-out”, “contra-front”, “contra-in”, and “ipsi-front”, respectively. Animals preferred to use their contralateral forepaw for contralateral targets, and their ipsilateral forepaw for the ipsilateral target (rare trials in which animals used the opposite forepaw were excluded). **a.** Trial-averaged movement trajectories of the contralateral forepaw (left) and ipsilateral forepaw (right) during the four-target reaching task. Trajectories are centered relative to the resting position of the paw (i.e. 500ms prior to touch). Colored squares indicate the latitude of the reach target. Trials where animals moved the wrong forepaw (i.e. not proximal to target) or both forepaws were excluded. **b.** Heatmaps showing normalized peri-movement activity of 61 SNr units during the four-target reach task. Units were sorted according to neural response latency during the “contra-front” target trials (for color scale see Fig. 8b). **c.** Normalized peri-reach activity ([−0.1, 0.2] s around reach onset) for 61 SNr units, during “contra-front” vs. “ipsi-front” target trials, where animals reached forward with the contralateral or ipsilateral paw, respectively. **d.** Normalized peri-reach activity ([−0.1, 0.2] s around reach onset) for 61 SNr units, during “contra-in” vs. “contra-out” target trials, where animals used the contralateral paw to reach inward or outward, respectively. In **c-d**, solid black dots indicate SNr units showing significant peri-movement responses in one or both movement types (permutation test; p<0.01; n=31/61, 19/61); red circles indicate SNr units whose peri-movement response amplitudes significantly differ across two movement types (permutation test; p<0.01; n=18/61, 7/61); grey circles represent the remaining unmodulated units.

**Figure 8—Figure Supplement 1.**
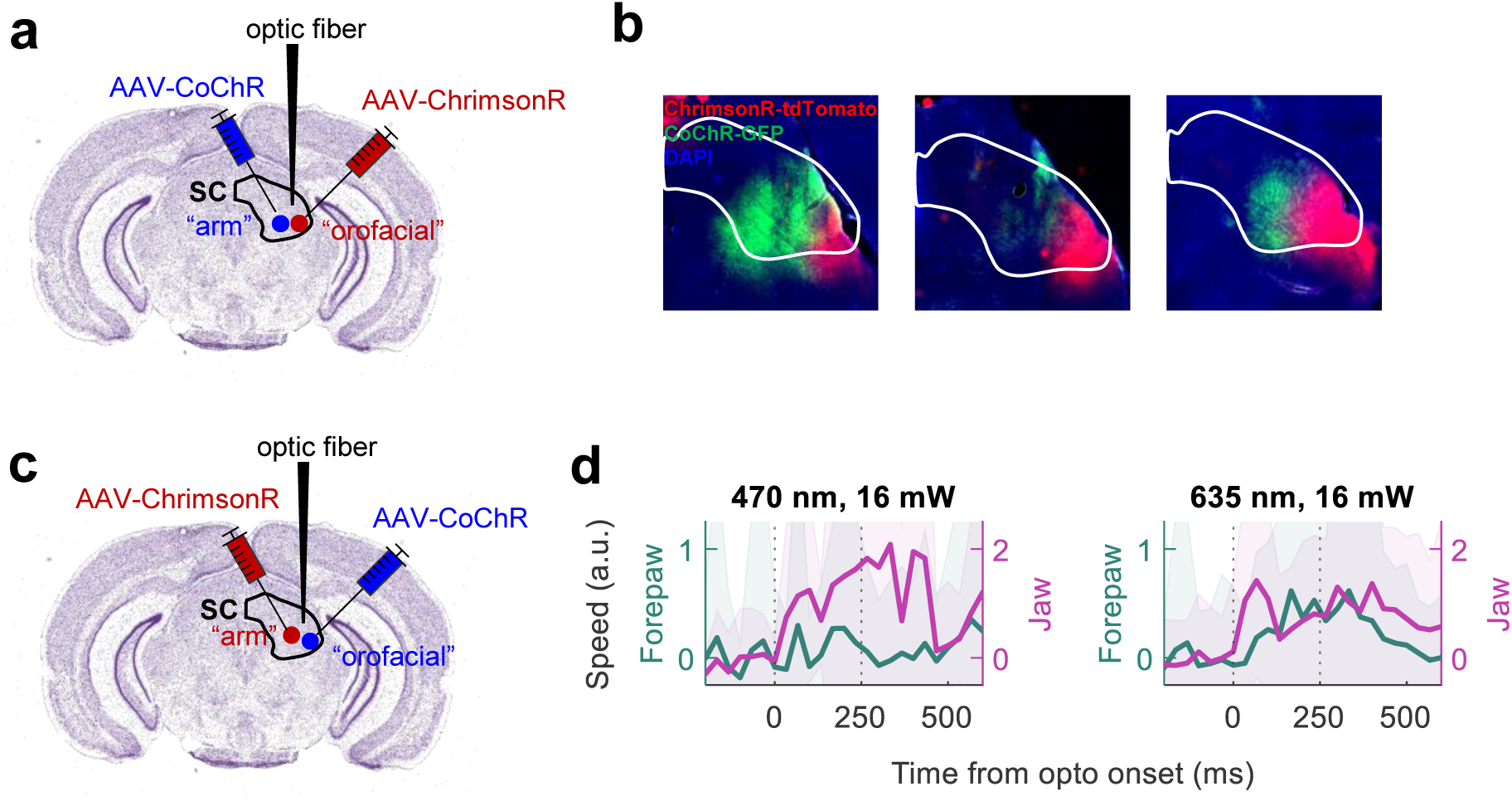
Verification of somatotopy in central vs. lateral SC. **a.** Schema of virus injection and optogenetic verification of arm vs. orofacial zones of motor SC. Adapted from the Allen Mouse Brain Atlas (2011). **b.** Histology showing the separation of viral expression in SC (3 animals). **c.** Schema of reversed viral injection. **d.** Optogenetically triggered movements of the forepaw and jaw by 470 nm vs. 593 nm lasers. Pulse duration = 250 ms.

## Notes

https://dataverse.harvard.edu/dataset.xhtml?persistentId=doi:10.7910/DVN/BZLOQS

### Competing Interest Statement

The authors have declared no competing interest.

### Summary of Updates

Added acknowledgement section; Updated formatting; Changed naming of supplemental figures from "Figure S1" to "Figure 1--Supplemental Figure 1"; Updated Methods section to include manufacturer address; Fixed some mislabeled figures references in the Methods section.

### Funding

- National Institute of Neurological Disorders and Stroke (NINDS): U19-NS113201
- National Eye Institute (NEI): P30-EY012196

